# Gravity-based microfiltration reveals unexpected prevalence of circulating tumor cell clusters in ovarian and colorectal cancer

**DOI:** 10.1101/773507

**Authors:** Anne Meunier, Javier Alejandro Hernández-Castro, Nicholas Chahley, Laudine Communal, Sara Kheireddine, Newsha Koushki, Nadia Davoudvandi, Sara Al Habyan, Benjamin Péant, Anthoula Lazaris, Andy Ng, Teodor Veres, Luke McCaffrey, Diane Provencher, Peter Metrakos, Anne-Marie Mes-Masson, David Juncker

## Abstract

Circulating tumor cells (CTCs) are rare (few cells per milliliter of blood) and mostly isolated as single cell CTCs (scCTCs). CTC clusters (cCTCs), even rarer, are of growing interest, notably because of their higher metastatic potential, but very difficult to isolate. Here, we introduce gravity-based microfiltration (GµF) for facile isolation of cCTCs while minimizing unwanted cluster disaggregation, with ∼85% capture efficiency. GµF from orthotopic ovarian cancer mouse models, from 17 epithelial ovarian cancer (EOC) with either localized or metastatic disease, and from 13 metastatic colorectal cancer liver metastasis (CRCLM) patients uncovered cCTCs in every case, with between 2-100+ cells. cCTCs represented between 5-30% of all CTC capture events, and 10-80% of CTCs were clustered; remarkably, in 10 patients, most CTCs were circulating not as scCTCs, but as cCTCs. GµF uncovered the unexpected prevalence and frequency of cCTCs including sometimes very large ones in EOC patients, and motivates additional studies to uncover their properties and role in disease progression.

## Introduction

Circulating tumor cells (CTCs) are shed from the tumor into the bloodstream, then circulate and reach distant locations where they can extravasate, proliferate and seed metastases. CTCs, found in the majority of epithelial cancers ^1^, represent a crucial intermediate in the metastatic cascade and their study has the potential to help improve patient care. CTCs are extremely rare (∼1-10 CTCs *vs*. < 10^6^ leukocytes and < 10^9^ erythrocytes per milliliter of blood) and display extensive molecular heterogeneity. CTCs in blood are correlated with poor outcome, recurrence and resistance to therapy ^1,2^. CTCs were first reported in 1869 ^3^, and interest has dramatically increased in the last decades following technical advances permitting simplified and effective isolation and quantification.

CTCs exist as single cells (scCTCs) and multicellular aggregates, called CTC clusters (cCTCs). scCTCs were traditionally isolated and identified based on the expression of the epithelial cell adhesion molecule (EpCAM) and cytokeratins (CK), but it became known that many scCTCs are disseminated following epithelial-mesenchymal transition (EMT) exhibiting downregulation of epithelial markers, and resulting in their enhanced motility and aggressiveness ^1^. It is believed that cCTCs do not arise from scCTC proliferation in blood but separate from the primary tumor as a cluster (collective dissemination) ^4^. Using mouse models, Aceto *et al*. found that cCTCs accounted for 3% of all captured CTC events, but have a 23- to 50-time higher metastatic potential than scCTCs ^4^. The presence of cCTCs in blood has been associated with worse outcome in patients with lung ^5^, breast ^6^, prostate ^7^, skin ^8^, bladder ^9^, pancreati^10^, head and neck ^11^, colorectal ^12-14^ and ovarian cancer ^1,4,15,16^.

The five-year survival in epithelial ovarian cancer (EOC) is ∼45% owing to late diagnosis and lack of effective therapy ^17^. In EOC, CTCs were detected in the blood of ∼10 to ∼60% patients ^18,19^, travelling as scCTCs ^20^ and rarely as cCTCs ^21^. EOC is also characterized by the formation of ascites within the peritoneal cavity, that contain tumor cells and provide a local microenvironment regulating the behavior of the tumor cells ^22^. In line with observations of Aceto *et al.* in blood, Al Habyan *et al.* ^23^ showed that disseminated tumor cells in ascites of EOC mouse models arise from spontaneous detachment as either single cells or clusters with cCTCs representing between 17-49% of all CTC capture events. Enriching cCTCs is therefore of great interest as their characterization could offer new insights into cancer dissemination, and help improve prognosis and treatments in cancer.

In colorectal cancer (CRC), CTC could often only be detected in a subset of patients. One study found ≥ 1 CTC per 7.5 mL of blood in 54%, and ≥ 3 CTC 7.5 per mL in 18.6 to 30% of patients prior to surgery^24^. In a more recent study, ≥ 1 CTC per 7.5 mL of blood were detected in 46% of pre-surgery CRC patients in all stages of the Union of International Cancer Control ^13^. The detection of one or more CTCs per 7.5 mL of blood is correlated to metastases, shorter progression-free and overall survival. High CTC count was associated with poor prognosis in metastatic CRC ^13,24-27^. cCTCs have only been rarely reported, while 1 to 5.4 cCTC per mL were detected in advanced and metastatic CRC ^28^.

Isolating cCTCs is more challenging given their rarity, their short lifespan, and their propensity to disaggregate under shear ^16^. Isolation technologies were initially developed and optimized for scCTCs, but following the occasional capture of cCTCs ^29-31^, further developments have been more efficient and selective at isolating clusters ^32,33^. Toner and colleagues pioneered two advances. One was based on deterministic lateral displacement tuned to deflects particles > 30 µm tested with isolated artificial breast cancer cell clusters spiked in blood ^34^. They also developed a filtration-based chip (Cluster-chip) using shifted triangular pillars forming 4000 parallel 12 × 100 μm^2^ slit openings designed to permit scCTC passage while capturing cCTCs ^16^. A more recent microfluidic chip-based device developed by the same group combined inertial focusing with repeated flow-shifting for cCTC isolation, enabling large volume blood samples processing capacity at a flow rate higher than 30 mL h^-1^ ^35^. The Parsortix system marketed by ANGLE ^36^ is a semi-automated system that flows under controlled pressure condition the blood sample through a microfluidic cassette. The sample flows through serpentine channels containing separation structures of increasing height laid out as steps, leading to a final critical gap (ranging from 10 µm to 4.5 µm) where larger cells (*e.g.* CTCs) are captured based on their size and deformability while red and white blood cells flow through. Parsortix has been adapted for cCTC capture, with a reported capture efficiency of > 97.2% for scCTC and > 99.3% for cCTC (cluster sizes not specified) spiked in blood ^37^.

Limitations of CTC isolation technologies include long processing times for cCTC isolation to reduce shear and the risk of cCTCs dissociation. For example, Parsortix requires ∼4 h to process 7.5 mL sample through a separation cassette with a 6.5 µm critical gap for large CTC capture. Cheng *et al*. employed an alternative approach using a three-dimensional scaffold to reduce shear stress, capturing both scCTCs and cCTCs at 50 μL min^-1^ (or 3 mL h^-1^), but aggregation and break-up on the filter could confound the quantification of cCTC filtration efficiency ^38^. Inertial focusing produces forces that are expected to cause cluster breakage, along with the disadvantage of losing small cells resulting in additional loss of information. Microfiltration is geometrically predisposed to low shear stresses because a small filter < 1 cm in diameter can accommodate hundreds of thousands of pores, each forming a parallel flow path with a very low flow and shear. Microfiltration of scCTC was first reported in 1964 ^39^ using track-etched membranes with a random pore distribution. However, to prevent pore overlapping, membrane porosity was limited to 3-5%, thus affecting flow rate and throughput ^40-43^. Isoporous membranes with regular pore pattern and high porosity fabricated by lithography can accommodate higher flow rates while maintaining low shear stress ^32,44,45^. We developed high performance polymer microfilters that satisfy key criteria for practical use, namely low-cost, durable, highly transparent, non-autofluorescent and highly porous (20-60%) ^46^. Following optimization, 8 mm-diameter microfilters with 8 μm pores, a 1:6 dilution of whole blood in phosphate buffer saline (PBS), and a flow rate of 0.1 mL min^-1^ set using programmable syringe pumps allowed us to recover > 80-95% of cancer cells spiked into blood ^47^.

Here, we introduce gravity-based microfiltration (GµF) for the efficient isolation of cCTCs using the in-house fabricated microfilters described above ^46^, and 3D printed cartridges ^47^. First, the gravity-flow of buffer, diluted and whole blood using filters with pores ranging from 8 to 28 μm in diameter was characterized, and flow rates, consistent with predictions, measured. The optimal flow rate and pore size for cCTCs isolation were determined using blood from healthy donors spiked with ovarian cancer cells from OV-90 cell line ^48^ comprising both single cells and clusters. Following a systematic characterization of the cluster morphology, the effect of shear on cluster disaggregation was characterized. Next, blood and ascites from ovarian orthotopic transplanted mice were processed and the number of scCTCs and cCTCs, along with the morphological features and the expression of molecular markers for aggressiveness and EMT were evaluated. Finally, to evaluate the clinical potential of GµF, blood samples from 30 cancer patients, including 17 EOC with advanced (metastatic) and localized disease (chemo-naïve and pre-surgery), and 13 colorectal cancer liver metastasis (CRCLM) patients were processed. The number and size of cCTC events and the proportion of scCTCs *vs*. cCTCs in the circulation were analyzed.

## Results

### Gravity-based microfiltration for selective isolation of clustered CTCs

Filter clogging by blood components is expected to increase the pressure and shear stress across the pores under constant flow rate conditions. We reasoned that increased shear would negatively affect the capture of clusters as it could contribute to disrupting or squeezing them through the pores. Thus we adopted gravity-based microfiltration (GµF), which provides constant pressure with constant total column heights (Htot) of 15, 20, and 30 cm (Figures 1A and S1) ^49^. As previously described ^47^, blood was diluted 1:6 in PBS and an 8 mm-diameter microfilter was clamped in the 3D printed cartridge (Figure 1B). Filters with pores of 8 μm diameter (Table S1) were used initially to characterize flow of GμF using PBS, diluted blood, and whole blood. The entire setup was assembled on a typical laboratory retort stand, with a footprint of 12 × 20 cm^2^, thus occupying little bench space with minimal hindrance to activities in the clinical laboratory.

**Figure 1.**
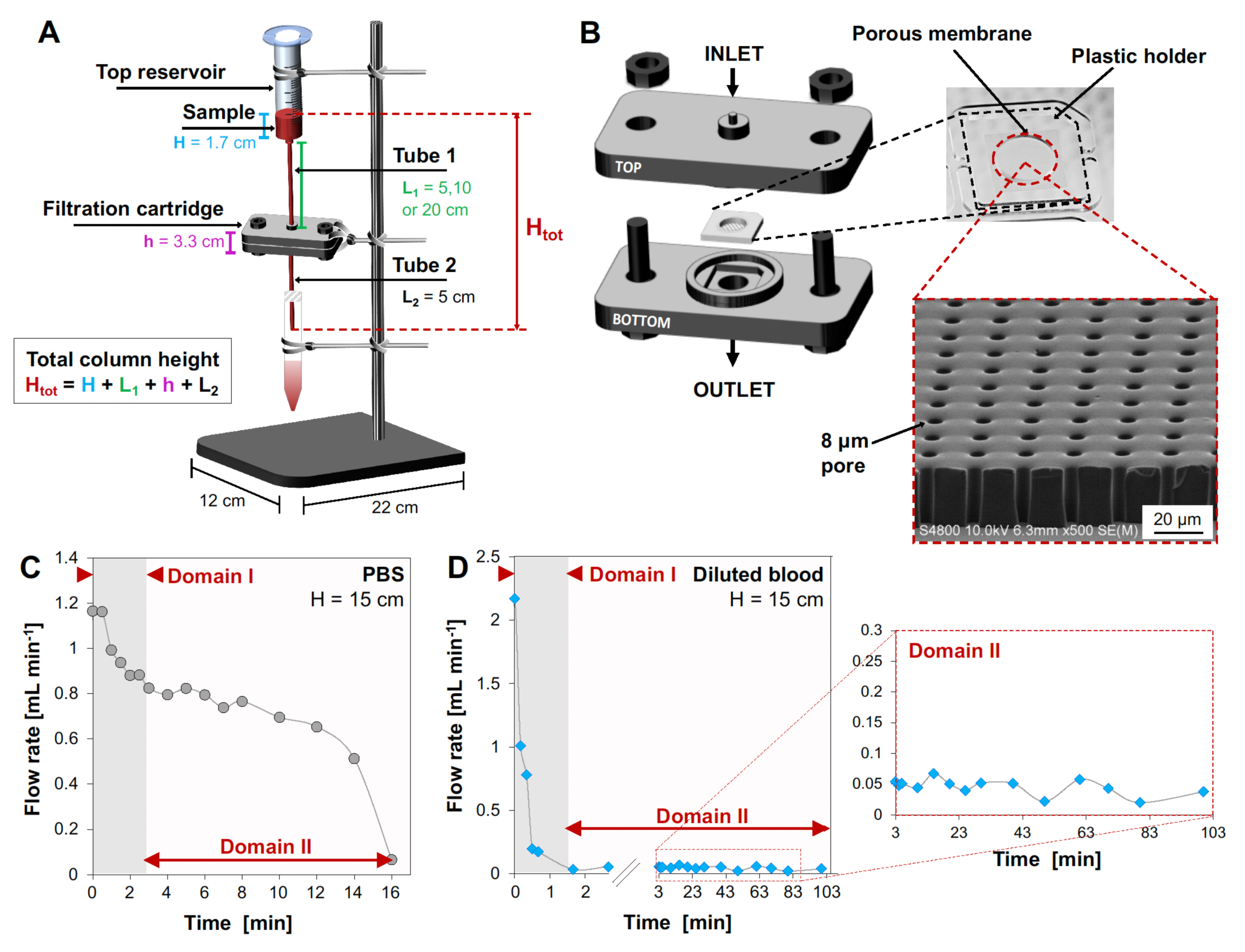
GµF for CTC enrichment. (A) Schematic of the GµF set-up. The total column height (Htot) determines the pressure and flow rate when considering the GµF flow resistance. (B) Exploded view of the filtration cartridge with filter, and close-ups of a polymer filter with 8 μm diameter pores. (C) and (D) Examples of time course of flow rate for 10 mL of (C) PBS and (D) blood diluted 1:6 in PBS filtered through an 8 μm filter with Htot = 15 cm, and close-up of domain II (pseudo-steady state) for diluted blood. See also Figures S1 and S2 and Tables S1 and S2.

For PBS-diluted and whole blood samples, the flow rate was highest at first and quickly decreased during the first seconds to minutes (termed Domain I). For PBS, the flow rate then decreased in concordance with the column height until the end of filtration, (termed Domain II, Figures 1C, S1B and Table S2). For diluted and whole blood, the flow rate then became constant which could last hours (also termed Domain II, Figures 1D and S1C-D). For example, for Htot = 20 cm (L1 = 10 cm), the flow rate in Domain II was ∼0.09 and ∼0.02 mL min^-1^ for diluted blood and whole blood, respectively. For both fluids, the flow rate in Domain II varied linearly with Htot. For diluted blood, similar behavior was observed for all filter pore sizes, and calibration curves (Figure S2) allowed the determination of the tube length to use for achieving a specific flow rate (Table S1).

We previously validated GµF for CTC isolation using two breast cancer cell lines (MDA-MB231 and MCF-7) and two kidney cancer cell lines (786-O and A-498) spiked in normal human blood ^47^. Here, to assess whether small differences between GµF and syringe pump driven flow were significant, diluted blood samples were spiked with ovarian cancer OV-90 single cells and clusters, divided into two equal volumes and filtered through 8 µm filters using both methods: one by GµF with a tube length L1 of 12 cm and 66 cm (nominal flow rates of ∼0.1 and ∼0.5 mL min^-1^, Table S1), and the other using a syringe pump with exactly 0.1 and 0.5 mL min^-1^. For 8-µm filters, the calculated shear stress (ρmax) was 0.58 Pa and 2.87 Pa and the transmembrane pressure (△P) was 5.75 Pa and 28.73 Pa, respectively (Table S3). After filtration, filters were rinsed with PBS, then cells were stained with DAPI for the nucleus, for epithelial cytokeratin (CK), and for CD45. This panel allowed discrimination of CTCs (CK^+^/CD45^-^/DAPI^+^) from white blood cells (WBCs) (CK^-^/CD45^+^/DAPI^+^) (Figure 2A). We did not observe significant clogging or biofouling of the device with flow remaining steady throughout the experiment.

**Figure 2.**
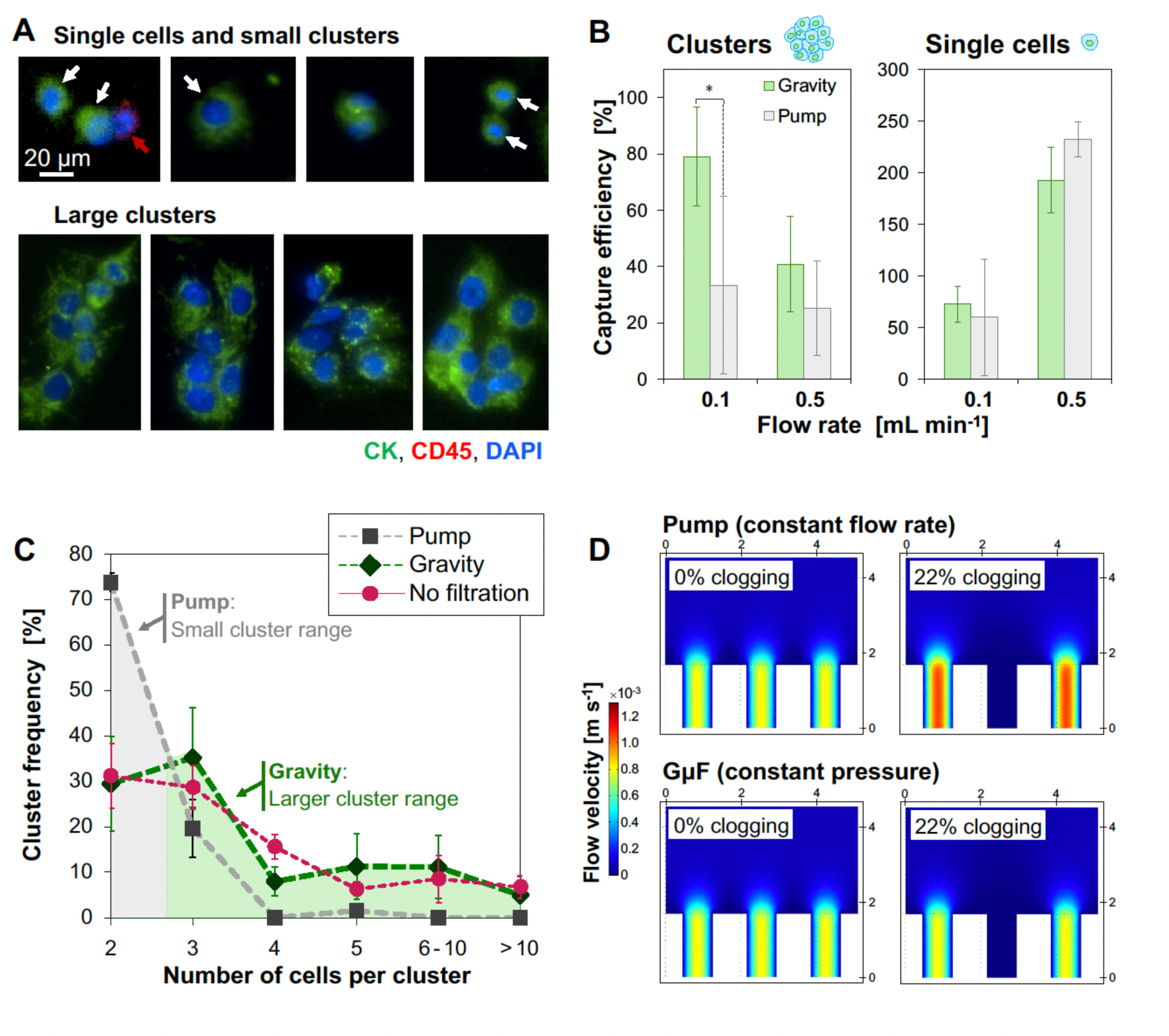
GµF captured larger clusters than syringe-pump driven flow. (A) Fluorescence images of OV-90 single cells and clusters captured using GµF and pump filtration using 8 μm filters. Cells were stained for cytokeratin (green) and CD45 (red). Nuclei were stained with DAPI (blue). Single cells (white arrows), small clusters and WBCs (red arrow) were captured using both configurations. Larger clusters, with > 5-6 cells were only captured with GµF. (B) Capture efficiency of clusters and single cells for GµF (green) and pump filtration (grey) at 0.1 and 0.5 mL min^-1^. (C) Size distribution of the clusters before filtration and captured using GµF and pump filtration at ∼0.1 mL min^-1^. For each replicate, a known number of OV-90 cells (∼120 clusters and ∼40 single cells) was spiked in diluted blood. Error bars correspond to the standard deviation of three replicated experiments. (D) Finite-element analysis comparing flow velocity in 8 μm pores for GμF and pump filtration at 0.1 mL min^-1^, for 0% and 22% clogging. See also Tables S1 and S3.

The capture efficiency for single cells and clusters showed opposing trends (Figure 2B). When the flow rate increased from 0.1 to 0.5 mL min^-1^, cluster capture efficiency decreased from ∼80% to ∼40% for GµF, and from ∼35% to ∼25% for pump-controlled flow, while single cell capture increased from 60% to 200% for GµF, and from 55% to 230% for pump filtration. Percentages over 100% are attributed to the break-up of clusters, which can account for the contradicting trends. These results highlight that GµF is more efficient at capturing clusters, in both cases, and that increasing the flow rate broke up 50% of clusters for GµF, and only ∼33% of clusters for syringe flow. This discrepancy may be explained by the fact that syringe flow already disrupted clusters at low flow rate.

Next, the size of OV-90 clusters after filtration at a nominal rate of 0.1 mL min^-1^ for GµF and syringe pump filtration was compared (Figure 2C). Consistent with cluster breakage, pump filtration showed more 2-cell clusters, but less 3-cell clusters, while 4-cell clusters and larger were essentially absent (∼1% of all capture events). Clusters of all sizes were captured by GµF, and 4-cell clusters and larger collectively accounted for ∼30% of all captured events.

Our previous work established 8 µm as the lower limit of pore size before clogging by WBCs becomes significant, and optimization showed that the number of clogged pores after filtration and rinsing was negligible (∼1%) ^47^. However, it is most likely that a significant number of WBCs is depleted during the rinse step. The proportion of clogged pore might be higher and vary during filtration. In GμF, although flow rate remained stable overall in Domain II, the slight fluctuations might reflect pore clogging, which would result in a slight pressure increase with constant flow rate pump filtration and explain the higher breaking rate. At 0.1 mL min^-1^, the maximum flow speed (ϖmax) through 8 µm pores is 826 µm s^-1^. For GμF, ϖmax remained constant irrespective of pore clogging but increased under constant-flow rate conditions during syringe pump filtration as shown by the FEA in Figure 2D. With 8 μm filter, for 22% clogging, ϖmax increased to ∼1058 µm s^-1^ and the resulting shear stress ι−max from 0.58 to 0.74 (Table S3).

### Capture of OV-90 cell clusters

Next, again using OV-90 cells, the optimal filter pore size for capturing the largest range of clusters was determined. Following spiking with ∼150 single cells and ∼100 clusters, each sample was divided into 6 equal aliquots and flowed through filters with 8 μm, 10 μm, 12 μm, 15 μm, 20 μm and 28 μm pores, respectively, using tube length L1 of 6 cm (flow rate ∼0.1 mL min^-1^) (Figure 3A and 3B). Single OV-90 cells were found on all filters and single cell capture efficiency strongly decreased with increasing pore size, while matching the size distribution of OV-90 cell before filtration, ranging between ∼6-50 μm in diameter. ∼70% of single cells were captured on 10 μm filters, dropping to ∼25% and 2% for 15 and 28 μm filters, respectively. Clusters were also found on all filters, and efficiency also decreased with pore size, but at a different rate and the efficiency was still ∼75% for 15 μm pores. Increasing pore size also yielded higher purity, with decreasing number of contaminating WBCs from ∼1000 to ∼200 WBCs following filtration of 1 mL using 8 μm and 15 μm filters, respectively ^47^.

**Figure 3.**
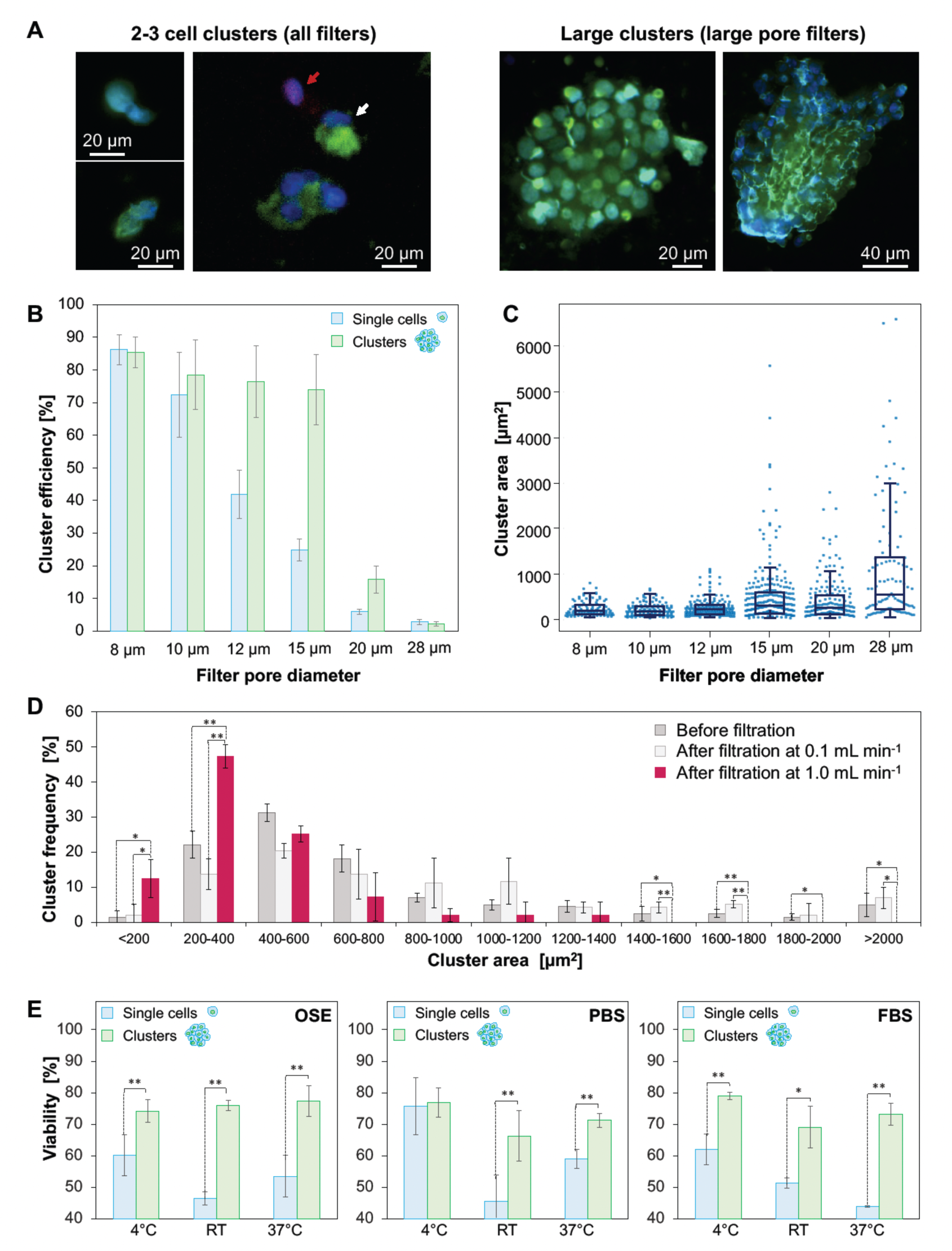
Characterization of clusters captured by GµF. (A) Fluorescence images of representative large and small clusters. Cells were stained for CK (green), CD45 (red) and with DAPI (blue). CTC-like cells (white arrow) are CK^+^/CD45^-^/DAPI^+^ and WBCs (red arrow) are CK^-^/CD45^+^/DAPI^+^. (B) Capture efficiency depending on pore size, measured with OV-90 single cells (blue) and clusters (green) spiked in diluted blood. (C) Scatterplot and box plot of the size distribution of clusters captured by serial GµF through filters with deceasing pore size (28, 20, 15, 12, 10 and 8 μm) for three replicated experiments (n = 145, 300 and 550 spiked clusters). The boxes range from 25^th^ and 75^th^ percentiles, the whiskers correspond to 91^st^ and 9^th^ percentiles, and the horizontal lines represent the medians. (D) Cluster size distribution before filtration (dark grey, n = 80, 39 and 37) and after filtration at 0.1 ml min^-1^ and release by flowing PBS at either at 0.1 mL min^-1^ (light grey, n = 61, 30 and 35) or 1 mL min^-1^ (red, n = 37, 31 and 19). (E) Viablility of single cells (blue) and clusters (green) after filtration, rinsing, and release using complete OSE culture medium, PBS, or FBS at 4°C, at RT (22-23°C), and after 5 h incubation in low adhesion plates (no processsing) at 37°C. Error bars correspond to the standard deviation of three replicated experiments. p < 0.01: **; p < 0.05: *. See also Figure S3.

Next, diluted blood samples spiked with OV-90 clusters were filtered successively through all filters by order of decreasing porosity and the cluster size distribution on each filter was determined by counting the number of cells they comprised and by measuring their surface area (Figure 3C and S3A). In this repeated filtration, all large clusters, from 6-100+ cells (> 1200 μm^2^, Øeq ∼40 μm), were captured on the three first filters with 28, 20 and 15 μm pores. As expected, 28 μm filters allowed for capturing the largest clusters with area ∼65000 μm^2^ (Øeq ∼90 μm) but most of the cluster population was captured on the following 20 μm and 15 μm filters, where cluster with area up to ∼2800 μm^2^ and ∼5500 μm^2^ were found, respectively. Small pore filters (8, 10, and 12 μm) mostly captured 2- and 3-cell clusters with areas ∼150- 300 μm^2^ (Øeq ∼14-19 μm), that could pass through the preceding larger pore filters. Small clusters were also found on larger pore size filters but to a lesser extent. Based on the absence of large clusters on downstream small pore filters, together with the very small area fraction of the filter covered by CTCs (typically << 0.1 %) and our previous work with single cells of other cell lines ^47^, we confidently exclude artefactual scCTC aggregation on the filter as a source of cCTCs during GµF. Interestingly, we observed that in small clusters (up to 5-6 cells), the diameter of the biggest cell increased with the pore size (Figure S3B). These results suggest that the capture efficiency of small clusters may not be dictated solely by the size of the cluster, but significantly depends on the size of its largest cell (and more accurately the largest nuclei) in agreement with Toner *et al*. who found that 20-cell clusters could traverse 5-10 μm constrictions in a microfluidic device following an unfolding process ^15^. The 15-μm filters contained the largest diversity of clusters, while no large clusters were found on the following 12, 10 or 8 μm filters, suggesting that 15 μm filters efficiently capture clusters, and were thus considered optimal. Using 0.1 mL min^-1^, 15-μm filters yielded △P of 1.34 Pa that resulted in a ϖmax of 338 μm s^-1^ and a ι−max of 0.13 Pa, below stress in capillaries ^50^ (Table S3), which provides further support that large clusters can be captured by GµF without disrupting them.

### Release of OV-90 clusters, cluster size distribution and viability

To characterize cell release from the filter, OV-90 single cells and clusters were first stained in suspension, then spiked in diluted blood. Samples were divided into three aliquots. One aliquot was used to determine clusters size distribution before filtration (positive control). The two other aliquots were filtered at 0.1 mL min^-1^, then cells were released by placing the cartridge upside down and flowing 5 mL of PBS in the reverse direction, either at 0.1 or 1.0 mL min^-1^, and their size distribution was characterized (Figure 3D).

The average area of clusters before filtration (795 ± 82 μm^2^) and after release at 0.1 mL min^-1^ (974 ± 57 μm^2^) was comparable, with clusters from ∼100 μm^2^ to 5000 and 10000 μm^2^, respectively. Following release at 1.0 mL min^-1^, average cluster area decreased to 486 ± 191 μm^2^, and the cluster distribution shrank with a strong increase in small cluster frequency and a significant loss of larger clusters. Clusters smaller than 400 μm^2^ represented 23 ± 6% of the total cluster population before filtration, 16 ± 7% after release at 0.1 mL min^-1^, and up to 59 ± 9% after release at 1 mL min^-1^. Although high flow rates are likely to help dislodging clusters, release efficiency decreased from ∼83% at 0.1 mL min^-1^ to ∼60% at 1.0 mL min^-1^, further highlighting the susceptibility of clusters to shear. At 0.1 mL min^-1^, GμF can both capture and release clusters while preserving their integrity.

Cell viability was measured after release from filters using blood samples spiked with ∼500 OV-90 single cells and clusters (Figure 3E). Dilution, rinsing, and release were performed with complete ovarian surface epithelial (OSE) culture medium, PBS, or fetal bovine serum (FBS) at 4°C and 23°C. A flow rate of 0.1 mL min^-1^, and filters with 15 μm and 8 μm pores were used to capture clusters and single cells, respectively. Single cells viability varied strongly with temperature; optimal conditions being PBS at 4°C (75.7 ± 9.1%). Cluster viability was independent of the tested parameters and stable at ∼70-80%, exceeding single cell viability in each case, except when using PBS at 4°C which raised single cell viability to the level of that of clusters. The preservation of cell-cell or cell-matrix interactions within clusters could explain their enhanced viability, and the lower effect of temperature compared to single cells. The loss of interaction with the extracellular matrix during shedding from the tumor induces activation of anoikis ^51^, and low temperature could slow down cell death mechanisms, thus accounting for increased single cell viability. As a positive control, cells were simply incubated for 5 h at 37°C in low-adhesion dishes and the viability was similar to that of cells released after GμF, indicating that neither single cell nor cluster viability were negatively affected by the GµF process. The full process from capture to release lasts ∼5 h, and cluster viability was ∼75%, and might be further increased by reducing filtration time by using filters with larger diameters.

### Capture of circulating tumor cell clusters (cCTCs) from mouse blood

GµF was tested with blood from ovarian orthotopic mouse models after injecting OV-90 or OVCAR-3 cells, with high and low invasive potential, respectively ^52^, and collecting their blood after sacrifice. Orthotopic xenograft results in spontaneous metastasis that closely mimic dissemination from the primary tumor in human cancers ^53^, thus recapitulating early events in disease progression. Half of each sample was saved for growth analysis, while the other half was filtered using 8 μm filters to capture the gamut of scCTCs and cCTCs. From the blood of three OV-90 mice, 505/5, 259/9 and 582/4 scCTCs/cCTCs were captured, cCTCs thus accounting for < 5% of the total CTC population, in agreement with published results ^4^. cCTCs with size between 2-100+ cells were found in every mouse.

Next, blood from a single OVCAR-3-GFP (green fluorescent protein) mouse was diluted and divided into four identical aliquots. One quarter was saved for further growth analysis, and the three others filtered through 15 μm filters at 0.1 mL min^-1^. Respectively 55, 40, and 49 cCTCs were found in each replicate, falling well within the variation expected based on Poisson statistics alone. The size of cCTCs again varied from 2-100+ cells, their frequency was highest for small clusters, with 2-cell and 3-cell clusters representing 27-33% and 5-12% of all cCTC events, respectively, and the larger clusters were rarer, with consistent results across the triplicates (Figure 4A and 4B). cCTCs as big as 30000-35000 μm^2^ (Øeq ∼0.6 mm) were captured directly from the mouse’s bloodstream, consistent with previous observations of large cCTC aggregates in blood ^15^. These results give us high confidence in the reproducibility and performance of GμF to isolate cCTCs spontaneously formed *in vivo*.

**Figure 4.**
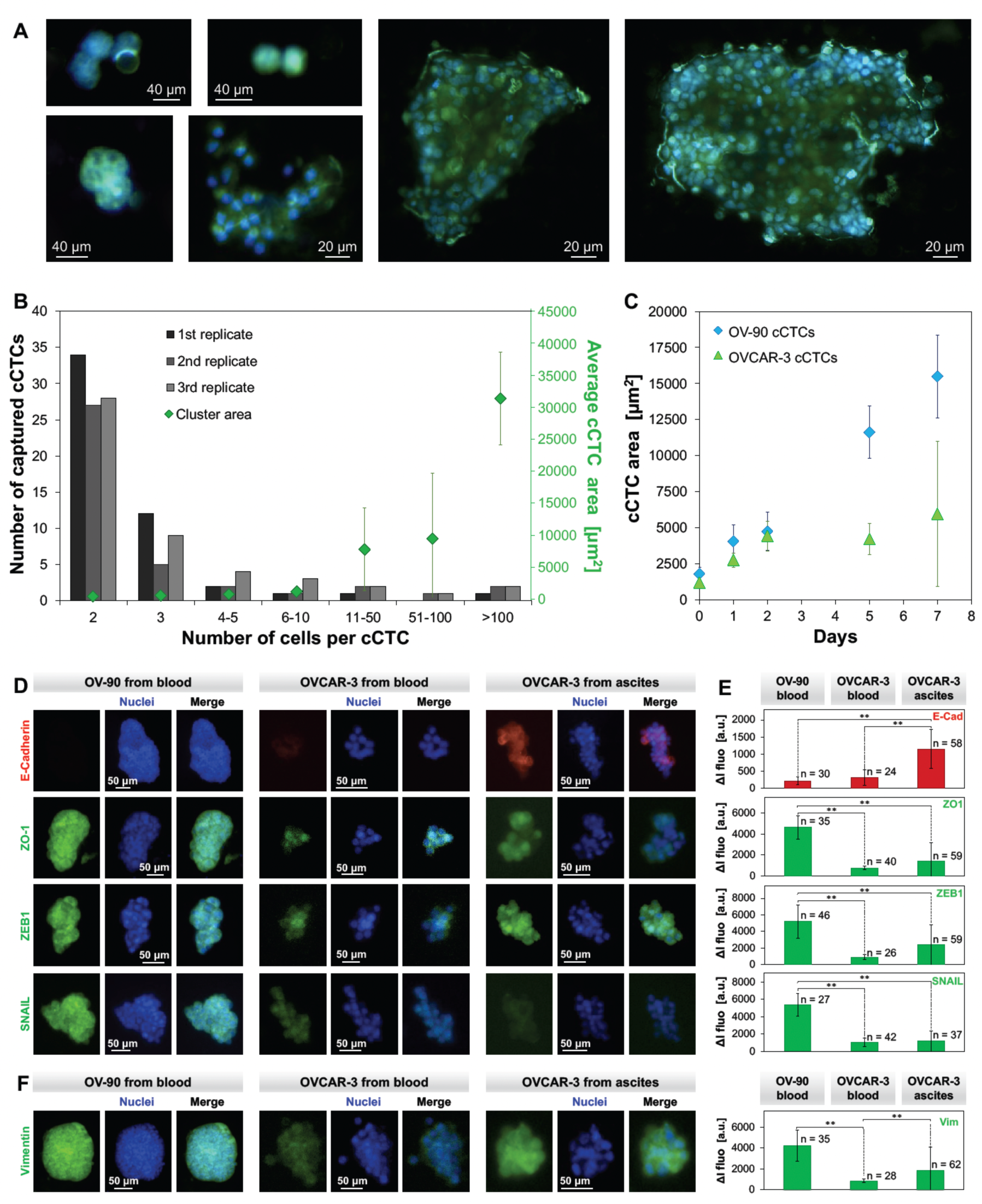
cCTCs captured from mouse. A. Examples of fluorescence images of OVCAR-3-GFP cCTCs captured from mouse blood. Nuclei were stained with DAPI. B. Size distribution of the OVCAR- 3-GFP cCTCs captured in three replicates from a single mouse. cCTC area is averaged over three replicates. C. Evolution of the average area of OV-90 and OVCAR-3-GFP cCTCs overtime during incubation in low adhesion culture flasks. Error bars correspond to the standard deviation of three replicated experiments. D. Fluorescence images of OV-90 cCTCs captured from mice blood, and OVCAR-3 cCTCs from the blood and ascites of the same mouse. cCTCs were stained for aggressiveness markers E-Cad, ZO-1, ZEB1 and Snail. Nuclei were stained with DAPI. E. Expression levels of E-Cad, ZO-1, ZEB-1 and Snail in OV-90 and OVCAR-3 cCTCs from blood or ascites with indication of statistical significance (p < 0.01: **). F. Additional staining for mesenchymal Vim. OVCAR-3 cCTCs from blood and ascites exhibit a hybrid E/M phenotype. Error bars correspond to the standard deviation between cCTCs. See also Figure S4.

The blood samples of OV-90 and OVCAR-3-GFP saved for growth analysis were filtered through 15 μm filters, and the cCTCs were released and cultured in regular and low-adhesion culture flasks at 37**°**C. Both OV-90 and OVCAR-3-GFP cCTCs adhered and spread after a few hours in adherent flasks forming confluent layers after 7-8 days. Migration assays revealed similar growth behaviors for both cell lines, where a full coverage of the cell-free area was reached after 6 days (Figure S4A-D). In low adhesion flasks, OV-90 and OVCAR-3-GFP cCTCs growth also occurred in suspension. After 7 days of incubation, the cluster average diameter increased from 1500-2000 μm^2^ (equivalent diameter of ∼40-50 μm) at day zero for both cell lines, to ∼6000 and ∼15500 μm^2^ for OVCAR-3-GFP and OV-90, respectively (Figure 4C). Both cell lines exhibited a broad cluster size distribution (Figure S4E-G) with cCTCs from 160 to more than 20000 μm^2^; OV-90 clusters were larger than OVCAR-3-GFP however, in agreement with a higher invasive phenotype. After day 2, OV-90 clusters kept increasing in size while OVCAR-3-GFP plateaued and at the same time a significant number of dying OVCAR-3-GFP was observed, likely due to a higher sensitivity of OVCAR-3 clusters to hypoxic conditions in the core.

### cCTCs can help understand dissemination in ovarian cancer

In EOC, Snail, a zinc finger transcription repressor, can activate zinc finger E-box-binding homeobox 1 (ZEB-1) that induce the loss of adhesion proteins such as E-cadherin (E-Cad) ^54^ and zonula occludens-1 (ZO-1) ^55^. As in many cancers, cell adhesion plays a critical role in the dissemination of EOC, and notably, the loss of E-Cad and ZO-1 ^55^ was correlated with disease aggressiveness by promoting tumor growth, invasiveness, and resistance to chemotherapy ^56,57^. cCTCs were isolated from the blood of two OV-90 and two OVCAR-3 mice, as well as from the ascites of one of the OVCAR-3 mice. After release, cCTCs were stained for aggressiveness markers: E-Cad, ZO-1, ZEB-1 and Snail (Figure 4D and 4E).

OV-90 cCTCs from blood strongly expressed ZO-1, ZEB-1 and Snail, but exhibited extremely low levels of E-Cad. The low E-Cad expression in mouse blood cCTCs contrasts the high expression level of E- Cad in OV-90 cell culture observed by Rosso *et al* ^58^. High E-Cad expression was also found in tumors of early stage (I-II) ovarian cancer patients, along with low expression of Snail and ZEB-1 that was consistent with their role as repressor of E-Cad. In contrast, in advanced stages (III-IV), E-Cad expression was low while Zeb-1 and Snail were highly expressed, suggesting repression of E-Cad. The observation in advanced ovarian cancer is in concordance with our mouse model results, which underwent spontaneous metastasis that closely mimics cell dissemination from the primary tumor at advance stage of the disease. The high level of mesenchymal marker Vim, along with low E-Cad expression in OV-90 cCTCs indicate a pronounced mesenchymal phenotype. This observation is consistent with the findings by Rosso *et al* where cell aggregates with a mesenchymal phenotype were able to survive under anchorage-independent conditions that are similar to CTC circulation in blood ^58^. Mesenchymal-like cell aggregates exhibit a high invasion potential that leads to metastasis, highlighting the aggressiveness of OV-90 CTCs. These observations support the contribution of EMT in cCTCs dissemination, and confirm the potential of GµF for molecular and functional analysis of living cCTCs.

To the best of our knowledge, cCTCs in blood and disseminated tumor cell clusters from ascites had not been compared, and interestingly, we found different phenotypes were dominant in each sample. OVCAR-3 cCTCs captured from blood displayed a weak expression of E-Cad and ZO-1, but also low expression of ZEB-1 and Snail, suggesting E-Cad expression is repressed by other mechanisms in this cell line. Likewise, OVCAR-3 clusters from ascites exhibited ZO-1, ZEB-1 and Snail expression levels that are in the same range or slightly higher than OCCAR-3 cCTCs from blood of the same mouse, and are significantly lower than those in OV-90 cells from blood. Interestingly, OVCAR-3 clusters from ascites had a much higher E-Cad expression than the other cCTCs (Fig. 4E). Gunay et al ^59^ observed very low expression of E-Cad in monolayer cell culture of OVCAR-3, but when cultured as spheroid, OVCAR-3 exhibited highly upregulated E-Cad expression. Spheroid formation within the peritoneal cavity is a hallmark of advanced stage ovarian cancer. The high E-Cad expression in OVCAR-3 clusters from mouse ascites is in agreement with Gunay’s observation in spheroids, which exhibited a loose aggregate morphology that is similar to the ascites OVCAR-3 clusters in advanced disease mouse models.

### Capture of cCTCs from cancer patients

To show the generality of GµF, we further analyzed CTCs from the blood of 17 EOC (Table 1), 13 CRCLM (Table 2) patients and 9 samples from healthy individuals. EOC patients included 10 high-grade serous carcinoma (OC1-OC10) with samples taken both before and after chemotherapy, and 7 other histological subtypes of EOC (endometrioid, clear-cell and mucinous carcinoma) with 5 of them at Stage IA to IC, and all taken before chemotherapy (OC11-15, and OC17). CRCLM patients (CR1-13) included 8 males and 5 females, with blood samples taken before surgery. 4 were treatment naïve and 9 received varied number of chemotherapy cycles.

**Table 1.** Epithelial ovarian cancer patient information. Cancer type, stage and grade at time of blood draw and CTC analysis as well as CA125 concentration in blood at the time of CTC analysis is provided. scCTC and cCTC counts are shown as number of events per mL.

| Patients | Cancer Type | Stage | Grade | Blood analysis | CA125<br>(U mL<sup>-1</sup>) | scCTC<br>(mL<sup>-1</sup>) | cCTC<br>(mL<sup>-1</sup>) |
| --- | --- | --- | --- | --- | --- | --- | --- |
| **High-grade serous carcinoma** |  |  |  |  |  |  |  |
| OC1 | Serous cystadenocarcinoma | IIIC | High grade | After surgery, pre-chemotherapy | 53 | 145 | 62 |
| OC2 | Serous cystadenocarcinoma | IIIC | High grade | After chemotherapy, cycle 4 | 98 | 82 | 3 |
| OC3 | Serous cystadenocarcinoma | IIIC | High grade | After chemotherapy, cycle 4 | 15 | 36 | 2 |
| OC4 | Serous cystadenocarcinoma | IIIC | High grade | After chemotherapy, cycle 4 | 52 | 22 | 1 |
| OC5 | Serous cystadenocarcinoma | IIIC | High grade | Post-chemotherapy, after cycle 6 | 10 | 59 | 5 |
| OC6 | Serous cystadenocarcinoma | IIIC | High grade | Pre-chemotherapy | 271 | 45 | 2 |
| OC7 | Serous cystadenocarcinoma | IVA | High grade | Pre-chemotherapy | 4354 | 68 | 9 |
| OC8 | Serous cystadenocarcinoma | IIIC | High grade | Pre-chemotherapy | 5153 | 123 | 39 |
| OC9 | Serous cystadenocarcinoma | IIIC | High grade | Pre-chemotherapy | 211 | 584 | 60 |
| OC10 | Serous cystadenocarcinoma | IVB | High grade | Pre-chemotherapy | 455 | 62 | 3 |
| **Other types of Epithelial Ovarian Cancer** |  |  |  |  |  |  |  |
| OC11 | Endometrioid carcinoma | IC | Low grade | Before surgery, pre-chemotherapy | 445 | 106 | 45 |
| OC12 | Clear-cell adenocarcinoma | IA | High grade | Before surgery, pre-chemotherapy | 21 | 155 | 18 |
| OC13 | Clear-cell adenocarcinoma | IC3 | High grade | At diagnosis | 706 | 74 | 8 |
| OC14 | Mucinous cystadenocarcinoma | IIIB | N/A | Before surgery, pre-chemotherapy | 171 | 25 | 3 |
| OC15 | Mucinous cystic tumor of borderline malignancy | IC1 | Borderline | Before surgery, pre-chemotherapy | 398 | 66 | 1 |
| OC16 | Ovarian Carcinoma | IIIB | Low grade | Pre-chemotherapy | 122 | 210 | 23 |
| OC17 | Endometrioid carcinoma | IC | Low grade | Pre-chemotherapy | 169 | 24 | 16 |
| **Time course analysis (Figure 6)** |  |  |  |  |  |  |  |
| OC1 | Serous cystadenocarcinoma | IIIC | High grade | After surgery, pre-chemotherapy (T<sub>0</sub>) | 53 | 146 | 62 |
|  |  |  |  | After chemotherapy, cycle 1 (T<sub>1</sub>) | 30 | 141 | 62 |
|  |  |  |  | After chemotherapy, cycle 2 (T<sub>2</sub>) | 14 | 132 | 4 |
|  |  |  |  | 8-month post-chemotherapy, cycle 6<br>(T<sub>3</sub>) | 11 | 11 | 2 |

**Table 2.** Colorectal Cancer Liver Metastasis patient information Sex, diagnosis, timing of blood draw, therapeutic treatment, as well as histopathological growth pattern (HGP) summary and operation room note when available. scCTC and cCTC counts are shown as number of events per mL.

| Patient | Sex | Blood Analysis | Previous Treatment | HGP | OR room notes | scCTC (mL<sup>-1</sup>) | cCTC (mL<sup>-1</sup>) |
| --- | --- | --- | --- | --- | --- | --- | --- |
| CR1 | F | Before surgery | 3 cycles chemotherapy | rHGP | 5 lesions: All at least 90% Replacement | 5 | 1 |
| CR2 | M | Before surgery | FOLFOX | rHGP | 1 lesion 100% replacement | 4 | 1 |
| CR3 | M | Before surgery | 7 cycles FOLFOX | rHGP | 1 lesion: Replacement | 24 | 9 |
| CR4 | M | Before surgery | 7 cycles CAPOX | rHGP | n/a | 35 | 12 |
| CR5 | M | Before surgery | Chemotherapy | rHGP | 1 lesion: 80% Replacement, 20% desmoplastic | 14 | 3 |
| CR6 | F | Before surgery | Naïve | rHGP | 2 lesions: 99% replacement, 1% desmoplastic | 17 | 4 |
| CR7 | F | Before surgery | Naïve | rHGP | 3 lesions: 2 (90% replacement, 10% pushing), 1 (98% replacement, 2% desmoplastic) | 26 | 2 |
| CR8 | M | Before surgery | 3 cycles CAPOX, then 1 cycle FOLFOX | dHGP | 1 lesion: 15% Replacement, 85% Desmoplastic | 56 | 1 |
| CR9 | F | Before surgery | 10 cycles FOLFOX, then 2 cycles FOLFIRI | dHGP | 3 lesions: Desmoplastic | 31 | 3 |
| CR10 | M | Before surgery | Naïve | dHGP | 1 lesion: Desmoplastic | 99 | 5 |
| CR11 | F | Before surgery | Naïve | pHGP | 1 lesion: 40% Replacement, 60% Pushing | 51 | 26 |
| CR12 | M | Before surgery | Chemotherapy | Mixed | 2 lesions: 1 pushing, 1 desmoplastic | 61 | 30 |
| CR13 | M | Before surgery | 2 cycles FOLFOX | Mixed | 4 lesions: 3 with more than 50% replacement, 1 lesion with more than 80% desmoplastic | 43 | 18 |

The blood samples were processed by filtering first with 15 μm followed by 8 μm filters to capture CTCs at 0.1 mL min^-1^. scCTCs and cCTCs were found in every patient blood sample, with different relative frequency for each patient. For EOC patients with high-grade serous carcinoma (Figure 5A), the number of scCTCs per mL of blood varied from 22 (OC4) to 584 (OC9), and the number of cCTCs from 1 (OC4) to 62 (OC1). For patients with good response after several cycles of chemotherapy, as indicated by low CA125 levels (Table 1), the proportion of cCTCs events was lower, varying from ∼4% (OC2) to ∼8% (OC5) of all CTC capture events. For OC1 (before chemotherapy, at T0), cCTCs represented up to ∼30% of all CTC capture events. For patients with other types of EOC at lower stage (Figure 5B), the number of scCTCs per mL of blood varied from 25 (OC14) to 210 (OC16), and the number of cCTCs per mL ranged from 1 (OC15) to 45 (OC11), and cCTCs represented from ∼1% (OC15) to ∼39% (OC17) of all CTC capture events.

**Figure 5.**
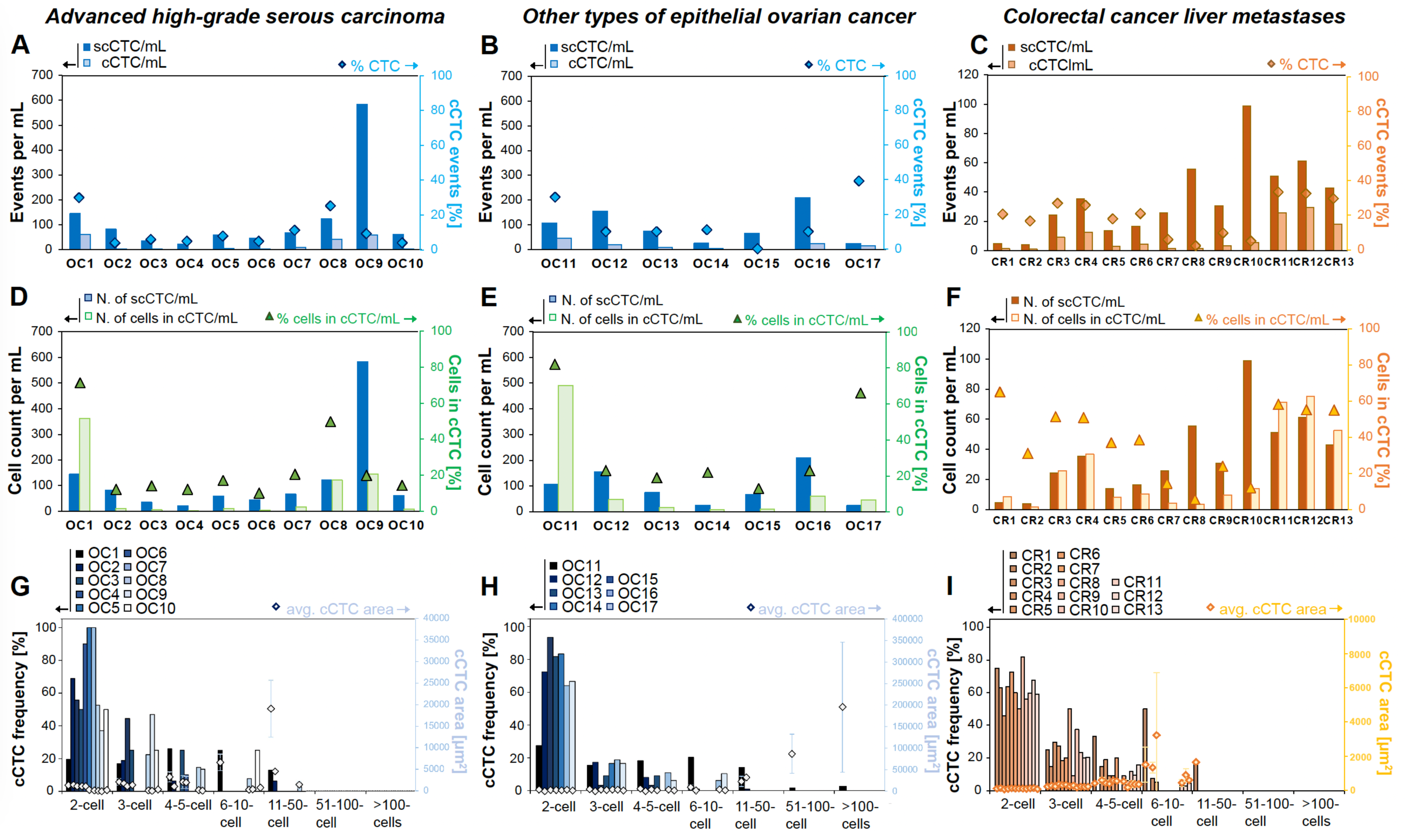
Count and morphological characterization of CTCs captured from the blood of EOC and CRCLM patients. (**A-C**) Concentration of scCTCs and cCTCs in the blood of patients OC1 to OC17 and CR1 to CR13. The diamonds represent the frequency of cCTC events. (**D-F**) Number of scCTCs and of cells in cCTCs per milliliter of blood, in each patient sample. The triangles correspond to the percentage of cells in cCTCs *vs.* all CTCs. (**G-I**) Size distribution of cCTCs captured from the blood of OC1 to OC17 and CR1 to CR13. The area covered by cCTC clusters is provided (dots) on an indicative basis. Error bars correspond to the standard deviation between cCTC within the cell number bin in each patient.

For CRCLM patients, the number of scCTCs per mL of blood varied from 4 (CR2) to 99 (CR10), and the number of cCTCs from 1 (CR1, 2, 8) to 30 (CR12). cCTCs were found in ≤ 10% of all CTC capture events for patients CR7-CR10, 17-27% in CR1-CR6, and ≥30% for CR11-13, with no apparent correlation with chemotherapy treatment. Note that the size of the tumor was not available.

Control experiments with healthy controls were performed to validate GμF processing and lend support to our clinical findings. Indeed, no cCTCs were detected in any healthy samples, and no scCTCs were detected in 6 out of 9; in three healthy controls, 1, 3 and 4 CK^+^/CD45^-^ single cells were detected in 3 mL of sample (*i.e.* ∼1 cell per mL). While the positive detection threshold for cancer is commonly set at ≥ 2 CK^+^/CD45^-^ scCTCs per 7.5 mL of blood ^60^, epithelial non-CTCs are commonly detected from the blood of healthy individuals at cell counts from 1.5 to 2 cells per mL^61,62^, and reaching as high as 21 cells per mL ^63^, and has been ascribed to shedding of epithelial cells into the bloodstream and/or blood collection device during venipuncture ^64^.

To better characterize the distribution of cCTCs and scCTCs, the number of cells captured as cCTCs was quantified by counting the number of cells in each cluster. The high transparency and low autofluorescence of the filters allowed us to precisely determine the number of cells for small clusters and estimate it for large clusters. In high-grade serous carcinoma (Figure 5D), cells in clusters represented up to ∼71% of all CTCs in the case of OC1, and was lower for patients having followed several cycles (4 to 6 cycles) of chemotherapy, varying from ∼12% (OC2) to ∼17% (OC5). For patients with low stage EOC (Figure 5E) and no chemotherapy, the proportion of cells in clusters varied from ∼13% (OC15) to ∼82% (OC11). Overall, in 11/17 EOC patients, cells in clusters corresponded to >15% of all CTCs, while they represented the majority of CTCs in 4/17 patients (OC1, 8, 11 and 66). For patients with CRCLM (Figure 5F), the proportion of cells in clusters varied from 6% (CR8) to 65% (CR1). Collectively, these results reveal an unexpected prevalence of cCTCs in patients with various histological subtypes of ovarian cancers at both high and low stage, and in colorectal cancer patients with liver metastasis.

In high-grade serous carcinoma, the size distribution of cCTCs varied from 2-50 cells (Figure 5G). In patients receiving therapy (OC1 at T1-T3, OC2-5) small cCTCs with 2 cells were the most frequent, corresponding to 50-90% of all cCTC events. 3-5 cells were found in 5/5 patients (10-50% of all cCTC events); and clusters with more than 11 cells were only found in OC1 and OC2. In patients who did not receive chemotherapy, small clusters with 2 cells were the most prevalent, corresponding to 53-100% of all cCTC events, except for OC1(at T0) and OC9 presenting 2-cell clusters in 19% and 37% of all cCTC events, respectively. In these two patients, clusters with 5 or fewer cells were still the most frequent prior to chemotherapy, representing 62% and 98% of all cCTC events, respectively. In patients with other EOC at lower stage and no chemotherapy, the size distribution of cCTCs varied from 2-100+ cells (Figure 5H), mirroring the one observed in mice. For OC11 and OC17, with endometrioid carcinoma, clusters of up to 3 cells represented 43% and 84% of all cCTCs, respectively. For OC11, clusters with more than 4 cells represented 57% of all cCTCs, including clusters with more than 50 and 100 cells. For OC12 and OC13 with clear-cell adenocarcinoma, OC14 and OC15 with mucinous disease, and OC16 with low-grade ovarian carcinoma, small clusters (2-3 cells) were the most frequent, representing 83-100% of all cCTCs events. No cluster larger than 3 cells were found in OC15 (borderline), and cluster larger than 6 cells were found in OC12, OC16 and OC17 and represented 2-10% of all cCTCs. CD45^+^/CK^-^ cells (*i.e.* WBC) counts were performed on 10 patient samples (OC1-5, and OC11-15) (Table S4). Of the 551 cCTCs isolated from the 10 samples, 18 were WBC^+^ and were found in 5/10 EOC patients, corresponding to 3.3% of all isolated cCTCs. The 18 WBC^+^ cCTCs comprised 3 to 5 clustered cells; 15 clusters had one WBC, two clusters had two WBC (OC3 and OC5), and one cluster had 3 WBCs (OC12).

All cCTCs in CRCLM patients were ≤ 10 cells, except for CR1 with 1 cCTC of > 10 cells (Figure 5I). In chemo-naïve patients, cCTCs of 2-3 cells were the most prevalent, in some cases corresponding to 79- 88% of all cCTC events (CR7, CR12 and CR13), except for CR1 who presents only cCTC of 4 cells or more. In CRCLM patients who underwent chemotherapy, cCTCs of 2-3 cells were also the most prevalent, representing 78%-100% of all cCTC events. There is no apparent correlation between cCTC size and chemotherapy cycles in this small cohort. WBCs were detected in the isolated cCTCs in 7/13 patients (Table S5). Of the 336 cCTCs isolated from the 13 samples, 75 were WBC^+^, corresponding to 22% of all cCTCs. Most WBC^+^ cCTCs were made up of 2 to 5 cells, while 8 WBC^+^ cCTCs were comprised of 6-10 cells, and contained no more than 4 WBCs. CK^+^/CD45^+^ cells were also observed in this work and by others ^65,66^. While these cells have been suggested to be hybrids deriving from the fusion of tumor cells and macrophages ^67^, further analysis is required to characterize and classify these atypical cells, they are thus not included in any counts in this work.

### Differential reduction of scCTC and cCTC during treatment of metastatic EOC high grade serous carcinoma patient

A time-course study was conducted for OC1 by collecting blood sample during the course of chemotherapy (Figure 6A). The concentration of CA125 in blood was used to track response to therapy and levels below the cut-off at 35 U mL^-1^ indicate a good response. The CA125 level of OC1 was reduced from >2000 U mL^-1^ at diagnosis to ∼50 U mL^-1^ after surgery and further decreased over the first three cycles of chemotherapy, plateauing at ∼10 U mL^-1^. CTCs analysis was performed at four times points: after surgery and before chemotherapy (T0), after the first cycle of chemotherapy (T1), a few days before the third cycle of chemotherapy (T2), and post treatment (T3); no pre-surgery samples were available for CTC isolation (Table 1). The number of scCTCs and cCTCs was stable between T0 and T1, with ∼140 scCTCs and ∼60 cCTCs per milliliter. Then, both scCTCs and cCTCs decreased in number as therapy progressed, but followed different trajectories. The number of scCTCs initially remained similar until the third cycle of chemotherapy, with 132 scCTCs per mL at T2, then decreased to 11 after 6 cycles of chemotherapy (T3). The number of cCTCs decreased from 62 to 2 in a gradual manner. Thus, while the CA125 level stopped changing after 3 cycles, the number of scCTCs and cCTCs continued dropping.

**Figure 6.**
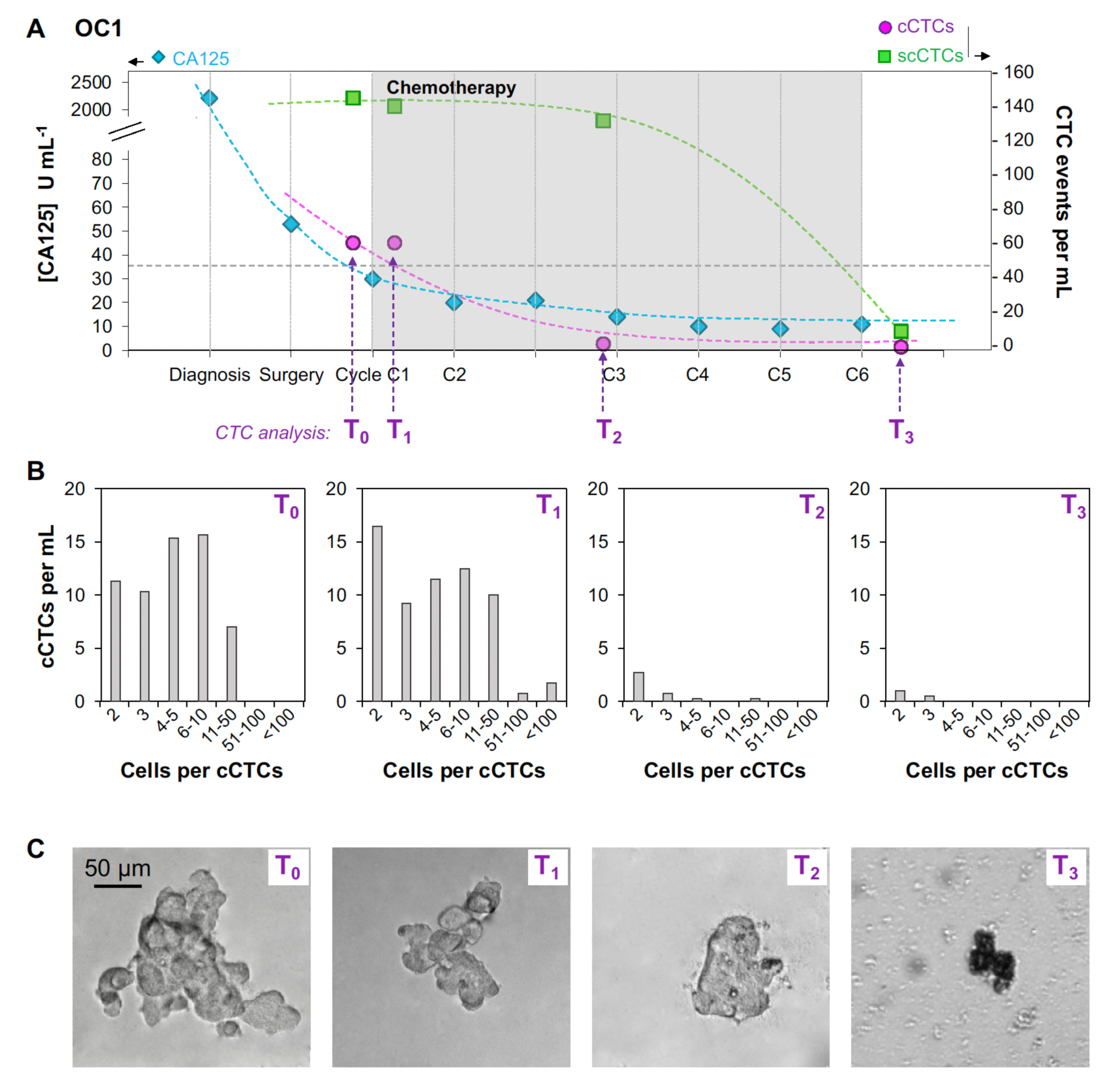
Time-course study for one metastatic EOC high grade serous carcinoma patient. **A.** Time course study for OC1 who responded to therapy, showing the CA125 concentration as well as the number of scCTC and cCTC events per mL of blood at different time points during chemotherapy. **B.** cCTCs size distribution and **C.** Representative bright field images of cCTCs captured from the blood of OC1 at four time points during follow-up.

Cluster size distribution was determined at the four time points (Figure 6B). Before chemotherapy (T0) and after the first cycle (T1), cCTC size distribution was similar, respectively with 36% and 42% of cCTCs with 2-3 cells, 52% and 38% of cCTCs with 4-10 cells, 12% and 16% of cCTCs with 11-50 cells. No cCTCs larger than 50 cells were captured at T0, and, at T1, clusters with 50-100+ cells corresponded to ∼4% of all cCTCs. This small increase in the number of large clusters between T0 and T1 is likely due to sampling variability as small blood volume were processed (3-4 mL), and larger clinical study would help determine its significance. Then, after the second cycle of chemotherapy (T2), in addition to the strong reduction in the number of cCTCs, the cCTCs size was significantly reduced, with the majority (∼94%) of cCTCs with 2-5 cells and only 1 cCTC with more than 11 cells. Finally, post-treatment (T3), only 2- and 3-cells clusters were captured. After the second cycle of chemotherapy (T2) and, to a much greater extent after the last cycle (T3), membrane blebbing and leakage of the cell contents was observed suggesting increased death of the cCTCs (Figure 6C). Together, these results indicate that chemotherapy may affect both the number and the size of cCTCs.

## Discussion

GμF using filters with 8 mm diameter and 15 μm pores for enriching cCTCs, and 8 μm pores for capturing the cCTCs that slipped through the large filters and scCTCs, is an effective method for capturing and selectively enriching the compendium of single cell CTCs and fragile clusters. Microfiltration was among the very first methods used to capture rare CTCs ^39,68^, and ‘rediscovered’ again ^44,45,69^, but typically only benchmarked for scCTC isolation. Two microfiltration methods, using ScreenCell and CellSieve filters, have recently been compared with CellSearch, and shown to be more efficient in detecting cCTCs. The higher efficiencies were mainly attributed to the marker-independent capturing approach ^70^. While clustered cell capture efficiencies of these two filtration methods were shown to range from 60-100%, they were determined using spiked-in mammospheres ^70^, which might have different properties than cCTCs. These methods also have shortcomings such as the need for leukocyte depletion to prevent clogging track-etched membrane (ScreenCell), and the requirement of a pump for photolithography fabricated membranes (CellSieve) ^70^. Cluster-Wells, a recent method developed for cCTC isolation, employs a 47 mm-diameter membrane etched with >100,000 microwells that was developed specifically for cCTC isolation. The geometry of the microwells retains clusters of 3 cells and more at nearly 100% efficiency, but at the expense of losing scCTCs ^71^. GµF efficiently captures both scCTCs and cCTCs, for which we ascribe the high capture yield to higher porosity filters (up to 40% porosity), gravity (pressure) driven flow, larger pores, and painstaking optimization.

GμF with 15 μm-pore filters and a flow rate of 0.1 mL min^-1^ (corresponding to a △P of 1.34 Pa, a ϖmax of 338 μm s^-1^, and a shear τ of 0.13 Pa) were found to be optimal for microfiltration in buffer. The velocity and shear in the pores is below the velocity (0.5-1.5 mm s^-1^) ^72^ and shear (0.5-2 Pa) ^50^ of blood capillaries. Unexpectedly, we achieved a higher cluster recovery rate with GμF *vs* constant-flow, at ∼80% *vs.* only ∼30%, respectively, with 8 μm filters while using the same nominal flow rate of 0.1 mL min^-1^ (Figure 2). We attributed the lower cluster recovery rate to higher cluster disruption under pump-driven flow even at low flow rate, which is consistent with the observation that clusters more than 4 cells were essentially absent using pump-driven filtration. To gain better understanding, we determined the number of single cells and clusters, and assessed the size distribution of OV-90 clusters spiked in blood, before and after filtration, which had not been studied previously to our knowledge. When the flow rate was 0.5 mL min^-1^, this analysis revealed twice as many single cells after filtration than were initially spiked into the sample, which we attributed to shear-driven cluster breakup. The cellular context bears significance in most cancers, and the shear forces in many commonly used isolation techniques are high enough to disrupt cCTCs. Hence, the analysis of the cluster number and size before and after filtration is necessary to assess the impact of a particular filtration method on cCTC breakup, and total counts of cCTCs and scCTCs. During detachment from 15-µm filters using flow reversal, higher flow rates increased the number of small clusters but reduced the apparent cluster release efficiency (∼60% at 1.0 mL min^-1^ *vs.*∼85% at 0.1 mL min^-1^), which again can be attributed to cluster break-up due to shear. A flow rate of 0.1 mL min^-1^ was thus deemed optimal for both capture and release of clusters with our setup.

The cluster capture efficiency for filters with pore sizes from 28 to 8 μm was characterized by sequentially filtering the same blood sample spiked with OV-90 clusters through filters with decreasing pore size and was found higher for pore sizes of 8-15 μm (Figure 3). The deformability and the large size distribution of both single cells and clusters limit the selectivity of size-based sorting. For example, small clusters squeezed through pores smaller than their nominal size, consistent with results by Toner and colleagues ^15^. With 15-μm filters, we recovered ∼75% of all clusters from blood, and ∼85% of captured clusters could be released, and confirmed that the cluster size distribution was the same before and after GμF.

To evaluate the potential for GμF for clinical use, 2.5-4 mL of blood from 30 patients, including 17 EOC (5 low stage and 12 high-grade serous carcinoma with metastasis) and 13 CRCLM were processed using 15-μm followed by 8-μm filters, allowing >85% recovery of both scCTCs and cCTCs. Whereas scCTCs are commonly isolated, and cCTCs have been observed with increasing frequency, they were presumed to be primarily associated with advanced metastatic diseases. Interestingly, Reduzzi *et al.* using ScreenCell microfilters detected cCTCs in 70% of early breast cancer patients versus only 20% in advanced, metastatic cases ^70^, whereas Boya *et al.* isolated cCTCs in 75% and 89% prostate and ovarian cancer patients using Cluster-Wells ^71^. In our study using GµF, both scCTCs and cCTCs were isolated from every patient whether with EOC or CRCLM, which for the former included patients with metastatic, localized, or borderline disease. Furthermore, samples from three castrate-resistant prostate and one renal cancer patients, all metastatic, were processed by GµF, and again, cCTCs found in every sample (as well as in the urine from the kidney cancer patient, results not shown). We did not detect any cell clusters in healthy controls. Taken together, these results suggest that (i) GµF is able to isolate cCTCs with high sensitivity, (ii) previous estimates of cCTC (and possibly of scCTCs) constitute a lower bound that is biased by technological limitations, and that (iii) cCTC may in fact be widely prevalent in a number of metastatic and non-metastatic cancers based on the observations made with breast cancer ^70^ and in this study EOC.

GµF using both 15-µm and 8-µm filters allowed to capture a broad range of CTCs from the blood of cancer patients. cCTCs from 2-100+ cells were captured, and small clusters, with 2-3 cells, were the most prevalent, representing from ∼43% to ∼100% of all clusters captured. EOC had the largest clusters, and both patient samples and mouse models found very large cCTCs occupying up to 350,000 μm^2^ on the microfilters, or ∼400 pores on a 15-μm filter, indicating that cCTCs can reach large sizes, consistent with other studies ^1,73,74^. cCTCs across a wider range were captured by GµF and with a higher yield than was previously reported for microfluidic systems such as the HB-Chip ^31^ and Cluster-Chip ^16^ that are only efficient in capturing small clusters from 4-12 cells. On the other hand, our result is in agreement with the detection of ovarian cancer cCTCs of >150 cells by Cluster-Wells ^71^. We also found that, in our EOC cohort, fewer WBC^+^ cCTCs were detected (3.3% of all cCTC events) than in the work by Boya *et al*. using Cluster-Wells (up to 26.4% in EOC cCTCs) ^71^, while the prevalence of WBC^+^ cCTCs from our CRCLM cohort (22%) is similar to the Boya et al.’s finding. Aceto *et al.* established the metastatic potential of cCTCs being 23- to 50-time the one of scCTCs based on the ratio of scCTCs *vs.* cCTCs captured using the HB-Chip from mouse blood. Given that the HB-Chip appears less effective than more recent methods such as GµF in cCTCs capture, the number of cCTCs may have been underestimated and by extension the metastatic potential of cCTCs, overestimated.

Studies on other commercialized platforms based on microfluidics (Parsortix) and microfiltration (CellSieve and ScreenCell) focused mainly on clinical translation. While these systems were reported to have close to 100% cCTC capture efficiencies in spike-in experiments, the size of the cCTCs were only estimated, and the number of cells in the spiked cCTCs were not determined ^37,70^. In this study the systematic analysis of the number of cells per clusters allowed us to account for the total number of CTCs circulating as cCTCs *vs.* scCTCs. Typically, ∼10-25% of CTCs were circulating as cCTCs, but in 10/30 patients most CTCs were not circulating as scCTCs, but as cCTCs, and at comparatively high concentration ∼45-60 cCTCs mL^-1^ in two patients.

In EOC, the initial response to therapy is generally good and can be traced by monitoring the CA125 drop in concentration in the blood. Relapse often occurs within 6-24 months, but we are currently lacking predictive blood biomarkers. In our small study, GμF revealed interesting differences among patients. The number of scCTCs, cCTCs, and the size of cCTCs were significantly lower for patients having followed several cycles of chemotherapy. Five patients (OC1-OC5) with serous cystadenocarcinoma (all metastatic) were analyzed. In one case (OC1) multiple blood samples were collected and processed before and during chemotherapy, and the time course of CTCs was compared to CA125. cCTC numbers were found to fall after the third chemotherapy cycle, mirroring CA125 concentration, but interestingly, the number of scCTCs remained stable and only dropped after the 6^th^ chemotherapy cycle. Thus, in line with previous observations supporting that CTC phenotype correlates with disease progression and recurrence in breast cancer ^74^, these results suggest that scCTCs and cCTCs could complement CA125 as an independent measure of response to therapy. The detection of CTCs in non-metastatic patients by GµF suggests also possible application to early diagnosis, which is supported by a study by Guo *et al.*, who found CTC counts to be more sensitive than CA125 to identify patients at high risk for ovarian cancer ^21^.

Limitations of this study include the small cohort size for both patients and controls. Further studies with larger numbers of patient and healthy control will be needed to confirm the unexpected prevalence of cCTCs found in the two cancers here, as well as studies with other cancers to evaluate the generality of these findings. CTC identification, counting and differentiation from other cell types are typically by manual and subjective immunofluorescence analysis of the stained cells and cell clusters ^76^, which is subject to interpersonal variation. The lack of CTC-specific markers also highlights the difficulty of using immunofluorescence for identifying all types of CTCs, especially given the heterogeneity of CTC. The implementation of CTC image recognition methods and machine learning algorithms could capture important image features, provide high-throughput CTC identification while reducing errors caused by manual interpretation, and would result in higher sensitivity and specificity.

In conclusion, optimized GμF is a simple yet powerful method for isolating scCTCs and notably cCTCs that are easily fragmented, but preserved thanks to gentle processing conditions. Spiked cCTCs were captured with ∼85% yield and released with 85% yield (combined yield 72%). cCTCs collected from patients and mouse models comprised between two to over one hundred cells. Our pilot study with 30 patients (17 EOC, including 5 non-metastatic ones, and 13 CRCLM), found cCTCs in every patient and in one third of the patients more cells were circulating as cCTCs than scCTCs. These results support the findings in recent studies where cCTCs were found to be prevalent in early breast cancer using low shear cCTC capture technologies ^70,75^, and challenge commonly held notions that (i) cCTCs are only found rarely, (ii) only in low numbers, and (iii) constitute only a small fraction relative to scCTCs.

Effective isolation of cCTCs along with scCTCs will allow studying their respective role in disease progression and metastasis, and their collective potential as surrogate biomarkers for diagnosis, prognosis and therapy monitoring. To allow comparison between different studies and methods, the capture yield for scCTCs and cCTCs, cluster break-up, size distribution and composition of cCTC will need to be characterized, and evaluated in studies with larger cohorts of patients and controls, and for different cancers.

## Acknowledgments

We acknowledge funding from the Canadian Institutes of Health Research (CIHR), as well as from the National Science and Engineering Research Council of Canada (NSERC) as part of the CHRP program (#446671-13). D.J. acknowledges support from Canada Research Chair (CRC). J.A.H.-C. thanks the Lloyd Carr-Harris Foundation and CONACyT for fellowships. L.M., FRQS research scholar, thanks the CIHR for financial support. A-M.M-M. and D.P. are researchers of the Centre de recherche du Centre hospitalier de l’Université de Montréal which receives financial support from the Fonds de recherche du Québec – Santé (FRQS). B.P. thanks the Institut TransMedTech for financial support. Tumor banking was supported by the Banque de tissus et données of the Réseau de recherche sur le cancer (RCC) of the Fond de recherche du Québec – Santé (FRQS), associated with the Canadian Tumor Repository Network (CTRNet). We would also like to acknowledge the MUHC Liver Disease Biobank for the CRCLM patient samples, supported through the MUHC Foundation. We gratefully acknowledge Dr. K. Rahimi from the department of pathology and cell biology at CRCHUM for advice and fruitful discussions. We thank Dr. Maïwenn Beaugrand, Adam Prager and Brittany Roberts for their help in experiments and data analysis.

## Author contributions

D.J. and A.M. designed the study. A.M. developed the GµF and characterized flow profiles. A.M. and S.K. optimized the GµF process using cell lines, J.A.H.-C. and N.C. fabricated microfilters and J.A.H.-C. conducted flow simulations. S.A.H., performed the orthotopic injection in mice. A.M. microfiltered mouse blood, while A.M., L.C., N.C. and N.D. processed patient samples. A.M., N.C., N.K., N.D. and B.P. conducted morphological and molecular characterization. D.P. and A-M.M-M. provided ovarian cancer cells and patient samples. A.L. and P. M. provided colorectal cancer liver metastasis patient samples. A.M., N.C., A.N. and D.J. analyzed the data. A.M., A.N. and D.J. wrote the manuscript. All authors discussed the results and commented and edited the manuscript.

## Methods

### Materials and reagents

All solutions were prepared using water from a Milli-Q system (resistivity: 18 MΩ cm; Millipore). Phosphate buffered saline (PBS, 1X, pH 7.4, Fisher Scientific), contains 11.9 10^-3^, 137.0 10^-3^ and 2.7 10^-3^ mol L^-1^ of phosphates (Na2HPO4 and KH2PO4), NaCl and KCl, respectively. Trypsin-EDTA, bovine serum albumin (BSA), bovine insulin and Tween 20 were obtained from Sigma-Aldrich. Triton X-100 and paraformaldehyde (PFA) were purchased from Fisher Scientific. OSE (Ovarian surface epithelial) medium, L-glutamine and HEPES were obtained from Wisent. RPMI (Roswell Park Memorial Institute) 1640 medium, fetal bovine serum (FBS) and 4’,6-diamidino-2- phenylindole (DAPI) were purchased from Life Technologies. Antibiotics (penicillin/streptomycin) were obtained from Invitrogen. Anti-human CD45-PE (cluster of differentiation 45, Cat. #FAB1430P), anti-CK 18-AF 488 (cytokeratin 18, labeled with Alexa Fluor 488, Cat. #IC7619G) and anti-E-Cad (E-cadherin, Cat. #MAB18382, from mouse) were obtained from R&D systems. Anti-Vim (Vimentin, Cat. #SAB4503081, from rabbit), anti-ZO-1 (Zonula occludens-1, Cat. #AB2272, from rabbit), anti-ZEB-1 (Zinc finger E-box-binding homeobox 1, Cat. #SAB3500514, from rabbit), and anti-Snail (Drosophila embryonic protein, Cat. #SAB2108482, from rabbit) were purchased from Sigma-Aldrich. Anti-EpCAM-PE (anti-epithelial cell adhesion molecule labeled with phycoerythrin, Cat. #12-9326-42), anti-c-MET-FITC (hepatocyte growth factor receptor, labeled with fluorescein isothiocyanate, Cat. #11-8858- 42), and detection antibodies goat anti-mouse-Alexa Fluor 647 (Cat. #A21240) and goat anti-rabbit-Cy3 (Cyanine 3, Cat. #A10520) were obtained from Fisher Scientific.

### Filter fabrication

The filter fabrication process, extensively described elsewhere ^46^, allows for the fabrication of porous membrane with pore diameters of 8, 10, 12, 15, 20 and 28 μm, referred to as X μm filters throughout the text. Briefly, pillar structures with a diameter corresponding to that of the pore to be created were prepared by standard photolithography and deep reactive-ion etching (DRIE). A UV-curable polymer cover coated on polyethylene terephthalate (PET) carrier was placed onto the pillars to close the structure. The formed cavity was then filled with Fluorolink® MD 700, which was then cured through UV exposure (2000-EC Series UV curing flood lamp, DYMAX). Finally, the blank cover was peeled off, and the molds were bathed in acetone for 15-20 min, for the membranes to self-de-mold from the pillars. Filters consist of a 20-40 μm-thick porous membrane heat bonded to a PMMA ring, which defines an 8 mm-diameter filter. Before filtration, filter surface was passivated by incubation in BSA (2% in PBS) to reduce non-specific adsorption on the filter.

### Filtration cartridge

The filtration cartridge (70 × 40 mm^2^) was designed with AutoCAD software (Autodesk Inc.) and 3D printed (Perfactory Micro EDU, Envision Tech) ^47^. It consists of a top (10 mm high) and bottom (15 mm high) parts with notches to place the filter between toric joints and allow for its alignment with the inlet and outlet. A silicone gasket and a pair of screws and bolts are used for sealing.

### Gravity-based microfiltration (GµF)

The GµF set-up consists of a 60 mL syringe (top reservoir) with its plunger removed, connected to the cartridge using a PEEK (Polyether ether ketone) tube (i.d. 0.75 mm, Sigma Aldrich). The overall setup is immobilized using a retort stand and the inlet tube is clamped before pouring the sample into the top reservoir. In this GµF, the height of the fluid column determines the pressure, and consequently the flow rate. Flow rate calibration was performed for whole and diluted blood (1:6 (v/v) in PBS), as well as for PBS for various filter porosities and tube lengths. After the first seconds to minutes of filtration, flow rate stabilized and fluctuate around a unique value for one to two hours. Average flow rates were measured by collecting sample droplets at known time intervals (from few seconds to minutes) during filtration. The average flow rate, measured in this stable regime was found to increase linearly with the initial sample column height. For each fluid, calibration curves of the average flow rate as a function of the sample height were established and used to adjust the tube length to achieve a specific flow rate (Figure S1 and Table S1).

### Cell culture

All culture media and solutions were sterile and filtered through a 0.2 μm filter. The ovarian cancer OV-90 cell line was developed in the laboratory of Drs. Provencher and Mes-Masson and has been well characterized ^48^. It was established from the cellular fraction of a patient’s ascites. OV-90 cells were maintained in OSE medium, supplemented with 10% FBS, 2% L-glutamine, 1% HEPES, and 1% (v/v) antibiotics (final concentrations of 100 I.U. mL^-1^ penicillin and 100 μg mL^-1^ streptomycin). Ovarian cancer OVCAR-3 cells were cultured in RPMI 1640 medium supplemented with 20% FBS, 1% (v/v) penicillin/streptomycin, and 0.01 mg mL^-1^ bovine insulin. Cell lines were validated by short tandem repeat (STR) profiling. For both cell lines, adherent cells were observed releasing single cells and clusters in their surrounding medium. The culture medium, containing released clusters, was changed every 1-2 days, and when adherent cells formed almost confluent layers (80-90%) in flasks, adherent cells were harvested using diluted trypsin. 200 μL of the cell suspension was re-suspended in 5 mL of culture medium in a new flask. All cell cultures were maintained in 5% CO2 at 37**°**C in 25 cm^2^ flasks (Corning).

### Spiking experiments with ovarian cell lines

The culture medium containing OV-90 clusters and very few single cells was harvested, centrifuged at 1300 rpm for 5 minutes, and cells and clusters resuspended in PBS. Single cells were obtained by trypsinization of the cell monolayers. Then, OV-90 single cells were diluted in PBS to obtain approximately 20-50 cells per microliter. In order to precisely determine the number of single cells and clusters, 10 μL droplets were placed between a microscope glass slide and a coverslip. The actual number of cells was manually counted twice on each slide and averaged on 10 droplets. Then, a known number of single cells and clusters was spiked in 1 mL blood samples (diluted 1:6 (v/v) in PBS). Blood was drawn from healthy volunteers (IRB study #BMB-08-012) into 10 mL CTAD tubes (citrate-based anticoagulant containing the platelet inhibitors theophylline, adenosine, and bipyridamole, BD Vacutainer). Samples were maintained at 4°C and processed within 72 h of collection.

### Flow velocity simulations

Modeling studies were performed using the COMSOL Multiphysics software. Flow velocity profiles were obtained by 3D simulations through a cell of 9 pores. Filter clogging was simulated through the same cells with 2/9 closed pores. For GμF (constant pressure), inlet pressure was fixed and for pump filtration a constant flow rate was applied. Additional details are provided in Table S3.

### Orthotopic mouse model of ovarian cancer

Female (8–12 weeks old) athymic nude mice (Crl:NU (NCr)-Foxn1nu; Charles River) were housed at the GCRC (Goodman Cancer Research Centre) animal facility and all procedures were conducted following ethics approval in accordance with the animal care guidelines at the Animal Resource Centre of McGill University. For orthotopic ovarian injections, 10 μL of Geltrex (Invitrogen) containing 7.5 × 10^5^ OV-90 or OVCAR-3 cells as a single-cell suspension were injected into the ovary. No leakage from the injection site was observed. When specified, and prior to the injection, OVCAR-3 cells were fluorescently labeled by lentiviral transduction (OVCAR-3-GFP). Lentivirus was produced in HEK293LT cells, by transfection with the lentiviral transfer vector plasmid (pWPI) that contains enhanced green fluorescent protein (eGFP) (Addgene plasmid # 12254), and provided by Dr. Didier Trono (Lausanne, Switzerland) ^77^. Ovarian tumors formed in ∼6 weeks and ascites were detected 8–10 weeks after injection. Animals that did not develop primary tumors, ascites, and metastases were excluded. Blood (∼0.5 mL) and ascites (∼0.5-2 mL) were collected under isoflurane anesthesia using a 23G needle and processed within 3-5 hours of collection.

### Epithelial ovarian cancer patients

Blood and tumor samples from EOC patients were collected with informed consent from the Centre Hospitalier de l’Université de Montréal (CHUM), in the Division of Gynecologic Oncology (Table 1). This part of the study involving human samples was approved by both institutional ethics committees: the Comité d’éthique de la recherche du CHUM (CÉR-CHUM) and the McGill research ethics office (IRB study #A05-M27-16B). Tumor stage was determined at time of surgery by a gynecologic oncologist. Histopathology and tumor grade were determined by a gynecological pathologist using criteria consistent with the International Federation of Gynecology and Obstetrics (FIGO) classification. Patient plasma CA125 levels were routinely measured during follow up. Blood samples used for CTC capture were kept at 4°C and processed within 1-14 hours of collection.

### Colorectal Cancer Liver Metastases blood samples

The study was done in accordance with the guidelines approved by McGill University Health Centre (MUHC) Institutional Review Board (IRB). Prior written informed consent was obtained from all subjects to participate in the study (protocol: SDR- 11-066). The study included a total of 13 CRCLM patients. Clinical data were collected for each patient through the locally established hospital database and medical records. Blood samples were collected fresh the day of the experiment in EDTA tubes and processed within 6 hrs.

### Blood samples from healthy individuals

9 healthy control samples were analyzed with approval by McGill research ethics office (study A04-M46-13B). Blood samples were collected in EDTA tubes and analyzed following the same procedure for the patient samples.

### CTC capture from blood sample

Before filtration, all samples (healthy and patient samples) were diluted 1:6 (v/v) in PBS. The protocol was revised with the addition of 0.05% of pluronic used for CRCLM patients. For filtration at constant flow rate using a syringe pump, samples were placed in a 10 mL syringe. For GµF, flow rate was adjusted by changing tube length based on calibration (Table S1). Initially, the inlet tube was clamped, and the sample was poured into the top reservoir, then filtration would start as the clamp was removed. Unless mentioned otherwise, filtration was performed at room temperature (22-23**°**C) and after filtration samples were rinsed twice with 5 mL of PBS at the same flow rate as that of filtration.

### Cell staining

Cell staining can be performed on the filter, directly in the cartridge after filtration as previously described ^47^ or after release in a culture dish. Cells were fixed with 3.7% paraformaldehyde (PFA), rinsed with PBS, permeabilized with 0.2% Triton X-100, then rinsed again with PBS. Blocking was performed with 1.0% BSA in PBS supplemented with 0.1% Tween 20. Then, for identification, cells were stained with anti-CK (cytokeratin) 18-Alexa Fluor 488 (2.0 μg mL^-1^) and anti-human CD45-PE (cluster of differentiation 45, labeled with phycoerythrin, 1.0 μg mL^-1^) to further distinguish cancer cells from blood cells in spiked, healthy control and clinical samples. For characterization, CTCs from mouse blood and ascites were stained using anti-E-Cad, anti-Vim, anti-ZO-1, anti-ZEB-1, or anti-Snail (1 mg mL^-1^), and GAM-647 or GAR-Cy3 (4.0 μg mL^-1^) for detection. Cells were rinsed with PBS, then their nucleus was counterstained with 4**′**,6-diamidino-2-phenylindole (DAPI, 0.1 μg mL**^−^**^1^).

### Cell release

Once filtration was over, the cartridge was placed upside down and 5 mL of PBS (or another fluid, where mentioned) were flown by gravity. Flow rate was controlled by adjusting the tube length. The fluid passes through the filter from the outlet to the inlet, thus mechanically detaching cells from the filter. The cell suspension was then centrifuged (1300 rpm, 5 min) and re-suspended in PBS before further staining or in culture medium for growth.

### Cell viability

Single cell and cluster viability was characterized after processing (dilution, filtration and release) with ovarian surface epithelial (OSE) culture medium (n=625/248, 690/626, and 508/186 single cells/clusters), phosphate buffer solution (PBS, n = 401/225, 199/58, and 474/238), or fetal bovine serum (FBS, n = 542/415, 384/184, and 459/308) using a live/dead kit (Thermofisher, #L3224). Cell suspensions were centrifuged, rinsed with PBS, centrifuged again and stained by incubation with 4.0 μmol L^-1^ of EthD-1 (red, dead cells) and 2.0 μmol L^-1^ of calcein AM (green, live cells) diluted in PBS for 45 min at room temperature. As a positive control, single cells and clusters, directly harvested from culture flasks, were seeded in ultra-low attachment well plates (Corning #3473) to avoid cell adhesion and then incubated for 5 hours at 37°C in OSE, PBS, or FBS. For negative control, dead cells were prepared by incubation in 70% methanol for 45 minutes. Viability was determined using fluorescence microscopy (excitation/emission wavelengths: 485/530 and 530/645 nm for calcein AM and EthD-1, respectively), corresponds to the ratio between the number of live single cells or clusters versus the total number of single cells or clusters per image, and was averaged over 5 images per condition and three replicated experiments.

### Fluorescence microscopy

Filters were placed upside down on the platform of an inverted microscope (TE-2000-E, Nikon) connected to a CCD camera (QuantEM 512SC, Photometrics), and fluorescence images were recorded with NIS-Elements Advanced Research software (Nikon), and analyzed with ImageJ (W. Rasband). Images were collected with a mercury arc lamp and using 41001 (blue, for AF 488, GFP, FITC and Cy3), 41004 (green, for PE and AF647), and 31000v2 (UV for DAPI) filter cubes (Chroma Technology Corp.). Cells are defined as CTC(-like) when they have a nucleus (DAPI) and express CK, a cytoplasmic protein of epithelial origin. WBCs also possess a nucleus but express CD45. For comparison and relative quantification of the expression levels of E-Cad, Vim, ZEB-1, ZO-1, Snail, EpCAM, and c-MET within different cluster models, same exposure time (1 s) was used for all images.

### Migration assay

Growth of OVCAR-3 and OV-90 clusters, isolated from mouse blood, was evaluated using migration assays. About 5×10^5^ cells mL^-1^ were seeded in each well of 2-well silicone inserts (Ibidi, Germany) placed on the bottom of a petri dish. Cells were incubated overnight, then the silicon insert was removed. The free-cell area was imaged and averaged over 10 images at different time points. The closure of the free-cell area overtime (% closure) was thus determined by comparison with the reference, measured right after removing the silicone inserts, and averaged over three replicated experiments (Figure S4A-D).

### Sphere forming assay

After capture from mouse blood, OV-90 and OVCAR-3 clusters were re-suspended in culture medium and incubated in ultra-low attachment wells. Spheres were grown for seven days, with gentle mixing once a day by pipetting. Samples were imaged right after seeding and over a few days of incubation using bright field microscopy. The area covered by clusters was measured using ImageJ software. Cluster proliferation in suspension was estimated by averaging the area of ∼400-500 clusters over a few days and for three replicated experiments (Figure 4C). Evolution of the area distribution of OV-90 and OVCAR-3 clusters over time is presented in Figure S4E-G.

### Statistical analysis

The data are presented as mean ± standard deviation measured over three replicates. Comparison of quantitative measures carried out on two independent groups was performed using unpaired two-tailed Student’s t-tests. Statistical significance was set as p < 0.05.

## Supplemental Information

Data generated or analyzed in this study are included in this published article and its supplementary information file.

## References

1 Pantel, K. & Speicher, M. R. The biology of circulating tumor cells. Oncogene 35, 1216–1224, doi:10.1038/onc.2015.192 (2016).

2 Poveda, A. et al. Circulating tumor cells predict progression free survival and overall survival in patients with relapsed/recurrent advanced ovarian cancer. Gynecol Oncol 122, 567–572, doi:10.1016/j.ygyno.2011.05.028 (2011).

3 Ashworth, T. A case of cancer in which cells similar to those in the tumours were seen in the blood after death. Med J Aust 14, 146–149 (1869).

4 Aceto, N. et al. Circulating tumor cell clusters are oligoclonal precursors of breast cancer metastasis. Cell 158, 1110–1122, doi:10.1016/j.cell.2014.07.013 (2014).

5 Gallo, M. et al. Clinical utility of circulating tumor cells in patients with non-small-cell lung cancer. Transl Lung Cancer Res 6, 486–498, doi:10.21037/tlcr.2017.05.07 (2017).

6 Costa, C. et al. Analysis of a Real-World Cohort of Metastatic Breast Cancer Patients Shows Circulating Tumor Cell Clusters (CTC-clusters) as Predictors of Patient Outcomes. Cancers (Basel) 12, doi:10.3390/cancers12051111 (2020).

7 Scher, H. I. et al. Circulating tumor cell biomarker panel as an individual-level surrogate for survival in metastatic castration-resistant prostate cancer. J Clin Oncol 33, 1348–1355, doi:10.1200/JCO.2014.55.3487 (2015).

8 Lucci, A. et al. Circulating Tumor Cells and Early Relapse in Node-positive Melanoma. Clin Cancer Res 26, 1886–1895, doi:10.1158/1078-0432.CCR-19-2670 (2020).

9 Gazzaniga, P. et al. Circulating tumor cells detection has independent prognostic impact in high-risk non-muscle invasive bladder cancer. Int J Cancer 135, 1978–1982, doi:10.1002/ijc.28830 (2014).

10 Effenberger, K. E. et al. Improved Risk Stratification by Circulating Tumor Cell Counts in Pancreatic Cancer. Clin Cancer Res 24, 2844–2850, doi:10.1158/1078-0432.CCR-18-0120 (2018).

11 Garrel, R. et al. Circulating Tumor Cells as a Prognostic Factor in Recurrent or Metastatic Head and Neck Squamous Cell Carcinoma: The CIRCUTEC Prospective Study. Clin Chem 65, 1267–1275, doi:10.1373/clinchem.2019.305904 (2019).

12 Yokobori, T. et al. Plastin3 is a novel marker for circulating tumor cells undergoing the epithelial-mesenchymal transition and is associated with colorectal cancer prognosis. Cancer Res 73, 2059–2069, doi:10.1158/0008-5472.CAN-12-0326 (2013).

13 Abdalla, T. S. A. et al. Prognostic value of preoperative circulating tumor cells counts in patients with UICC stage I-IV colorectal cancer. PLoS One 16, e0252897, doi:10.1371/journal.pone.0252897 (2021).

14 Heidrich, I., Abdalla, T. S. A., Reeh, M. & Pantel, K. Clinical Applications of Circulating Tumor Cells and Circulating Tumor DNA as a Liquid Biopsy Marker in Colorectal Cancer. Cancers (Basel) 13, doi:10.3390/cancers13184500 (2021).

15 Au, S. H. et al. Clusters of circulating tumor cells traverse capillary-sized vessels. Proc Natl Acad Sci USA 113, 4947–4952, doi:10.1073/pnas.1524448113 (2016).

16 Sarioglu, A. F. et al. A microfluidic device for label-free, physical capture of circulating tumor cell clusters. Nat Methods 12, 685–691, doi:10.1038/nmeth.3404 (2015).

17 Lheureux, S., Gourley, C., Vergote, I. & Oza, A. M. Epithelial ovarian cancer. Lancet 393, 1240–1253, doi:10.1016/S0140-6736(18)32552-2 (2019).

18 Judson, P. L. et al. Preoperative detection of peripherally circulating cancer cells and its prognostic significance in ovarian cancer. Gynecol Oncol 91, 389–394., doi:10.1016/j.ygyno.2003.08.004 (2003).

19 Fan, T., Zhao, Q., Chen, J. J., Chen, W.-T. & Pearl, M. L. Clinical significance of circulating tumor cells detected by an invasion assay in peripheral blood of patients with ovarian cancer. Gynecol Oncol 112, 185–191, doi:10.1016/j.ygyno.2008.09.021 (2009).

20 Allard, W. J. et al. Tumor cells circulate in the peripheral blood of all major carcinomas but not in healthy subjects or patients with nonmalignant diseases. Clin. Cancer Res. 10, 6897–6904 (2004).

21 Guo, Y.-X. et al. Diagnostic value of HE4+ circulating tumor cells in patients with suspicious ovarian cancer. Oncotarget 9, 7522–7533, doi:10.18632/oncotarget.23943 (2018).

22 Kim, S., Kim, B. & Song, Y. S. Ascites modulates cancer cell behavior, contributing to tumor heterogeneity in ovarian cancer. Cancer Sci 107, 1173–1178, doi:10.1111/cas.12987 (2016).

23 Al Habyan, S., Kalos, C., Szymborski, J. & McCaffrey, L. Multicellular detachment generates metastatic spheroids during intra-abdominal dissemination in epithelial ovarian cancer. Oncogene 37, 5127–5135, doi:10.1038/s41388-018-0317-x (2018).

24 Cohen, S. J. et al. Relationship of circulating tumor cells to tumor response, progression-free survival, and overall survival in patients with metastatic colorectal cancer. J Clin Oncol 26, 3213–3221, doi:10.1200/JCO.2007.15.8923 (2008).

25 Arrazubi, V. et al. Circulating Tumor Cells in Patients Undergoing Resection of Colorectal Cancer Liver Metastases. Clinical Utility for Long-Term Outcome: A Prospective Trial. Ann Surg Oncol 26, 2805–2811, doi:10.1245/s10434-019-07503-8 (2019).

26 Bidard, F. C. et al. Circulating Tumor Cells and Circulating Tumor DNA Detection in Potentially Resectable Metastatic Colorectal Cancer: A Prospective Ancillary Study to the Unicancer Prodige-14 Trial. Cells 8, doi:10.3390/cells8060516 (2019).

27 Silva, V. S. E. et al. Baseline and Kinetic Circulating Tumor Cell Counts Are Prognostic Factors in a Prospective Study of Metastatic Colorectal Cancer. Diagnostics (Basel) 11, 502, doi:10.3390/diagnostics11030502 (2021).

28 Zhang, D. et al. Circulating tumor microemboli (CTM) and vimentin+ circulating tumor cells (CTCs) detected by a size-based platform predict worse prognosis in advanced colorectal cancer patients during chemotherapy. Cancer Cell Int 17, 6, doi:10.1186/s12935-016-0373-7 (2017).

29 Brandt, B. et al. Isolation of prostate-derived single cells and cell clusters from human peripheral blood. Cancer Res. 56, 4556–4561 (1996).

30 Reddy, R. M. et al. Pulmonary venous blood sampling significantly increases the yield of circulating tumor cells in early-stage lung cancer. J Thorac Cardiovasc Surg 151, 852–858, doi:10.1016/j.jtcvs.2015.09.126 (2016).

31 Stott, S. L. et al. Isolation of circulating tumor cells using a microvortex-generating herringbone-chip. Proc Natl Acad Sci USA 107, 18392–18397, doi:10.1073/pnas.1012539107 (2010).

32 Ferreira, M. M., Ramani, V. C. & Jeffrey, S. S. Circulating tumor cell technologies. Mol Oncol 10, 374–394, doi:10.1016/j.molonc.2016.01.007 (2016).

33 Rushton, A. J., Nteliopoulos, G., Shaw, J. A. & Coombes, R. C. A Review of Circulating Tumour Cell Enrichment Technologies. Cancers (Basel) 13, doi:10.3390/cancers13050970 (2021).

34 Au, S. H. et al. Microfluidic isolation of circulating tumor cell clusters by size and asymmetry. Sci Rep 7, 2433, doi:10.1038/s41598-017-01150-3 (2017).

35 Edd, J. F. et al. Microfluidic concentration and separation of circulating tumor cell clusters from large blood volumes. Lab Chip 20, 558–567, doi:10.1039/c9lc01122f (2020).

36 Miller, M. C., Robinson, P. S., Wagner, C. & O’Shannessy, D. J. The Parsortix Cell Separation System-A versatile liquid biopsy platform. Cytom A 93, 1234–1239, doi:10.1002/cyto.a.23571 (2018).

37 Gkountela, S. et al. Circulating Tumor Cell Clustering Shapes DNA Methylation to Enable Metastasis Seeding. Cell 176, 98–112 e114, doi:10.1016/j.cell.2018.11.046 (2019).

38 Cheng, S.-B. et al. Three-dimensional scaffold chip with thermosensitive coating for capture and reversible release of individual and cluster of circulating tumor cells. Anal Chem 89, 7924–7932, doi:10.1021/acs.analchem.7b00905 (2017).

39 Seal, S. H. A sieve for the isolation of cancer cells and other large cells from the blood. Cancer 17, 637–642, doi:10.1002/1097-0142(196405)17:5<637::aid-cncr2820170512>3.0.co;2-i (1964).

40 Vona, G. et al. Isolation by Size of Epithelial Tumor Cells. Am J Pathol 156, 57–63, doi:10.1016/s0002-9440(10)64706-2 (2000).

41 Desitter, I. et al. A new device for rapid isolation by size and characterization of rare circulating tumor cells. Anticancer Res 31, 427–441 (2011).

42 Coumans, F. A., van Dalum, G., Beck, M. & Terstappen, L. W. Filtration parameters influencing circulating tumor cell enrichment from whole blood. PLoS One 8, e61774, doi:10.1371/journal.pone.0061774 (2013).

43 Netto, G. J. & Schrijver, I. Genomic applications in pathology. (Springer, 2015).

44 Adams, D. L. et al. The systematic study of circulating tumor cell isolation using lithographic microfilters. RSC Adv 4, 4334–4342, doi:10.1039/C3RA46839A (2014).

45 Lim, L. S. et al. Microsieve lab-chip device for rapid enumeration and fluorescence in situ hybridization of circulating tumor cells. Lab Chip 12, 4388–4396, doi:10.1039/c2lc20750h (2012).

46 Hernandez-Castro, J. A., Li, K., Meunier, A., Juncker, D. & Veres, T. Fabrication of large-area polymer microfilter membranes and their application for particle and cell enrichment. Lab Chip 17, 1960–1969, doi:10.1039/C6LC01525E (2017).

47 Meunier, A. et al. Combination of Mechanical and Molecular Filtration for Enhanced Enrichment of Circulating Tumor Cells. Anal Chem 88, 8510–8517, doi:10.1021/acs.analchem.6b01324 (2016).

48 Provencher, D. et al. Characterization of four novel epithelial ovarian cancer cell lines. In Vitro Cell Dev Biol - Anim 36, 357–361, doi:10.1290/1071-2690(2000)036<0357:COFNEO>2.0.CO;2 (2000).

49 Ripperger, S., Gösele, W., Alt, C. & Loewe, T. in Ullmann’s Encyclopedia of Industrial Chemistry (Wiley-VCH Verlag GmbH & Co. KGaA, 2012).

50 Ballermann, B. J., Dardik, A., Eng, E. & Liu, A. Shear stress and the endothelium. Kidney Int. 54, S100–S108, doi:10.1046/j.1523-1755.1998.06720.x (1998).

51 Frisch, S. M. & Francis, H. Disruption of epithelial cell-matrix interactions induces apoptosis. J Cell Biol 124, 619–626, doi:10.1083/jcb.124.4.619 (1994).

52 Lokman, N. A., Elder, A. S., Ricciardelli, C. & Oehler, M. K. Chick chorioallantoic membrane (CAM) assay as an in vivo model to study the effect of newly identified molecules on ovarian cancer invasion and metastasis. Int J Mol Sci 13, 9959–9970, doi:10.3390/ijms13089959 (2012).

53 Lengyel, E. et al. Epithelial ovarian cancer experimental models. Oncogene 33, 3619–3633, doi:10.1038/onc.2013.321 (2014).

54 Cano, A. et al. The transcription factor Snail controls epithelial-mesenchymal transitions by repressing E-cadherin expression. Nat Cell Biol 2, 76–83, doi:10.1038/35000025 (2000).

55 Polette, M. et al. β-Catenin and ZO-1: shuttle molecules involved in tumor invasion-associated epithelial-mesenchymal transition processes. Cells Tissues Organs 185, 61–65, doi:10.1159/000101304 (2007).

56 Haslehurst, A. M. et al. EMT transcription factors snail and slug directly contribute to cisplatin resistance in ovarian cancer. BMC Cancer 12, 91, doi:10.1186/1471-2407-12-91 (2012).

57 Wang, Y.-L. et al. Snail promotes epithelial-mesenchymal transition and invasiveness in human ovarian cancer cells. Int J Clin Exp 8, 7388–7393 (2015).

58 Rosso, M. et al. E-cadherin: A determinant molecule associated with ovarian cancer progression, dissemination and aggressiveness. PLoS One 12, e0184439, doi:10.1371/journal.pone.0184439 (2017).

59 Gunay, G. et al. The effects of size and shape of the ovarian cancer spheroids on the drug resistance and migration. Gynecol Oncol 159, 563–572, doi:10.1016/j.ygyno.2020.09.002 (2020).

60 Allard, W. J. et al. Tumor cells circulate in the peripheral blood of all major carcinomas but not in healthy subjects or patients with nonmalignant diseases. Clin Cancer Res 10, 6897–6904, doi:Doi 10.1158/1078-0432.Ccr-04-0378 (2004).

61 Sun, Y. et al. CTC phenotyping for a preoperative assessment of tumor metastasis and overall survival of pancreatic ductal adenocarcinoma patients. EBioMedicine 46, 133–149, doi:10.1016/j.ebiom.2019.07.044 (2019).

62 Green, B. J. et al. Phenotypic Profiling of Circulating Tumor Cells in Metastatic Prostate Cancer Patients Using Nanoparticle-Mediated Ranking. Anal Chem 91, 9348–9355, doi:10.1021/acs.analchem.9b01697 (2019).

63 Tsai, W. S. et al. Circulating tumor cell enumeration for improved screening and disease detection of patients with colorectal cancer. Biomed J 44, S190–S200, doi:10.1016/j.bj.2020.09.006 (2021).

64 Castle, J., Morris, K., Pritchard, S. & Kirwan, C. C. Challenges in enumeration of CTCs in breast cancer using techniques independent of cytokeratin expression. PLoS One 12, e0175647, doi:10.1371/journal.pone.0175647 (2017).

65 Lustberg, M. B. et al. Heterogeneous atypical cell populations are present in blood of metastatic breast cancer patients. Breast Cancer Res 16, R23, doi:10.1186/bcr3622 (2014).

66 Reduzzi, C. et al. Abstract P2-26-06: Association between CK+/CD45+ circulating tumor cells (CTCs) and circulating tumor DNA (ctDNA) alterations in advanced breast cancer patients. Cancer Research 83, P2–26-06-P22-26-06, doi:10.1158/1538-7445.Sabcs22-p2-26-06 (2023).

67 Clawson, G. A. et al. “Stealth dissemination” of macrophage-tumor cell fusions cultured from blood of patients with pancreatic ductal adenocarcinoma. PLoS One 12, e0184451, doi:10.1371/journal.pone.0184451 (2017).

68 Fleischer, R. L., Price, P. B. & Symes, E. M. Novel filter for biological materials. Science 143, 249–250 (1964).

69 Tang, Y. et al. Microfluidic device with integrated microfilter of conical-shaped holes for high efficiency and high purity capture of circulating tumor cells. Sci Rep 4, 6052, doi:10.1038/srep06052 (2014).

70 Reduzzi, C. et al. Circulating Tumor Cell Clusters Are Frequently Detected in Women with Early-Stage Breast Cancer. Cancers (Basel) 13, 2356–2375, doi:10.3390/cancers13102356 (2021).

71 Boya, M. et al. High throughput, label-free isolation of circulating tumor cell clusters in meshed microwells. Nat Commun 13, 3385, doi:10.1038/s41467-022-31009-9 (2022).

72 Arfors, K.-E., Bergqvist, D., Intaglietta, M. & Westergren, B. Measurements of blood flow velocity in the microcirculation. Ups J Med Sci 80, 27–33, doi:10.3109/03009737509178987 (1975).

73 Lecharpentier, A. et al. Detection of circulating tumour cells with a hybrid (epithelial/mesenchymal) phenotype in patients with metastatic non-small cell lung cancer. Br J Cancer 105, 1338–1341, doi:10.1038/bjc.2011.405 (2011).

74 Yu, M. et al. Circulating breast tumor cells exhibit dynamic changes in epithelial and mesenchymal composition. Science 339, 580–584, doi:10.1126/science.1228522 (2013).

75 Krol, I. et al. Detection of clustered circulating tumour cells in early breast cancer. Br J Cancer 125, 23–27, doi:10.1038/s41416-021-01327-8 (2021).

76 Hong, B. & Zu, Y. Detecting circulating tumor cells: current challenges and new trends. Theranostics 3, 377–394, doi:10.7150/thno.5195 (2013).

77 McCaffrey, L. M., Montalbano, J., Mihai, C. & Macara, I. G. Loss of the Par3 polarity protein promotes breast tumorigenesis and metastasis. Cancer Cell 22, 601–614, doi:10.1016/j.ccr.2012.10.003 (2012).

