## Supplemental File for "Gravity-based microfiltration reveals unexpected prevalence of circulating tumor cell clusters in ovarian and colorectal cancer"

### Table of Contents

**A. Gravity-based microfiltration (GµF)**

**Figure S1.** Schematic of the GµF set-up and gravity flow characterization.

**Table S1.** Filter characteristics and geometry of the GµF set up.

**Table S2.** Flow resistance in the GµF set-up.

**Figure S2.** GµF calibration for all pore sizes.

**Table S3.** Flow characteristics in the GµF set-up.

**B. Cluster area and cell diameter**

**Figure S3.** Cluster area and cluster's cells diameter.

**C. Growth of OV-90 and OVCAR-3 clusters captured from mouse blood**

**Figure S4.** Growth of OV-90 and OVCAR-3-GFP clusters previously isolated from mouse blood.

**D. WBCs in cCTCs isolated from EOC patient samples**

**Table S4.** WBC count in cCTCs isolated from 10 EOC patient samples.

**E. WBCs in cCTCs isolated from CRCLM patient samples**

**Table S5.** WBC count in cCTCs isolated from 13 CRCLM patient samples.

### A. Gravity-based microfiltration (GµF)

#### GµF set-up

The GµF setup (Figure S1A) consists of a 60 mL syringe (top reservoir, i.d. = 26.7 m) with its plunger removed, connected to the cartridge using a PEEK (Polyether ether ketone) tube (Tube 1, i.d. = 0.75 mm, $L_1$ = 5, 10 or 20 cm, Sigma Aldrich). The cartridge outlet is connected to a second tube (Tube 2, i.d. = 0.75 mm, $L_2$ = 5 cm). Both tubes are connected to the cartridges using Luer-Lock connector fittings (top and bottom junctions). Within the cartridge, the microfilter is clamped between a pair of o'rings, leaving an 8 mm diameter filtration area with pore diameter from 8 to 20 µm. Filter characteristics are provided in Table S1. The overall setup is immobilized using a retort stand and the inlet tube is clamped before pouring the sample into the top reservoir.

In gravity-based filtration, the height of the fluid column determines the pressure, and consequently the flow rate. Here, the total height of the fluid column ($H_{tot}$) includes the sample height (H) in the top reservoir, the tubes length ($L_1$ and $L_2$) and the thickness of the filtration cartridge. Thus, modification of the tube length ($L_1$) will allow for changing flow rate.

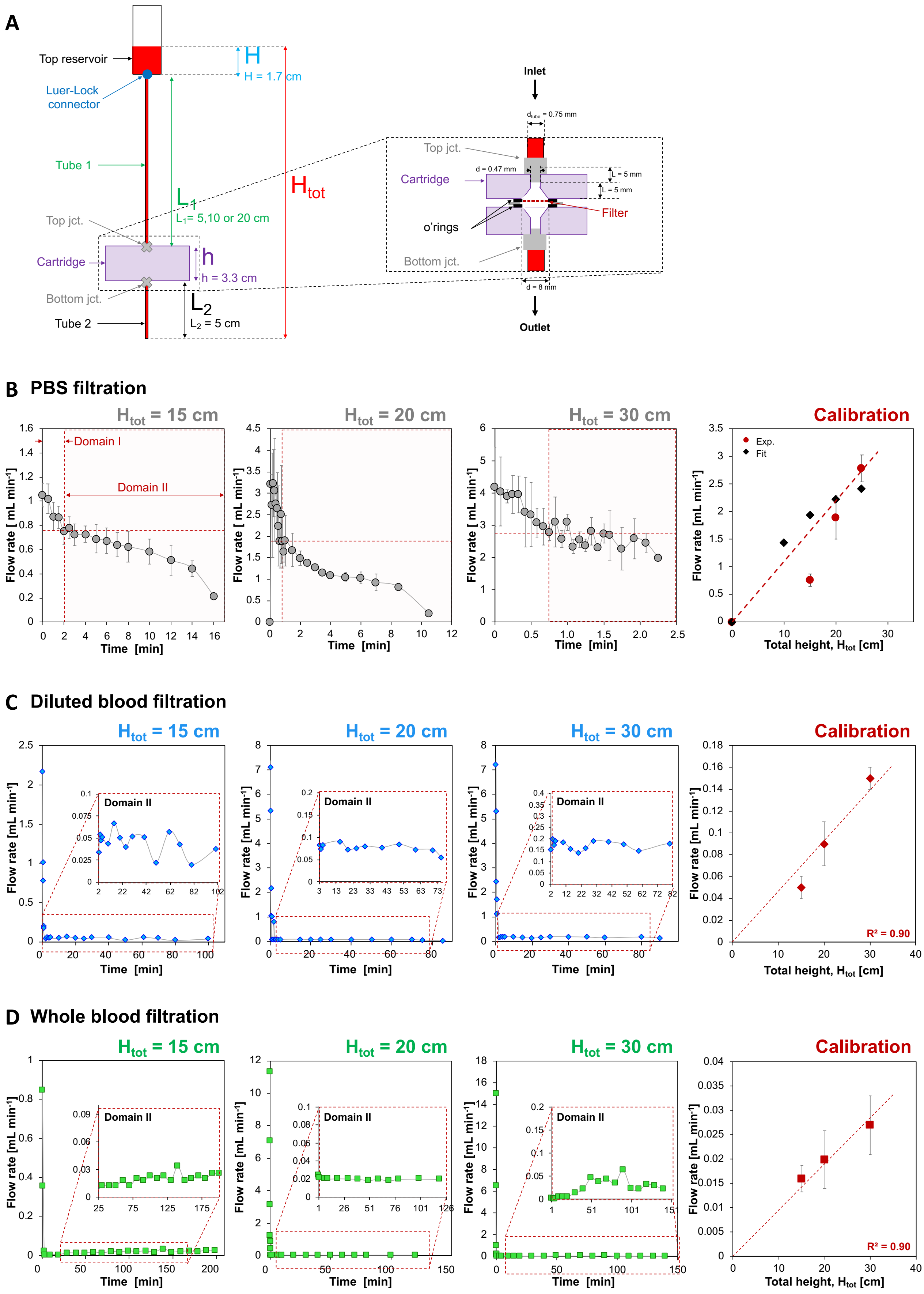

A
Top reservoir
Luer-Lock connector
Tube 1
Top jct.
Cartridge
Bottom jct.
Tube 2
H
H = 1.7 cm
L1
L1= 5,10 or 20 cm
h
h = 3.3 cm
L2
L2 = 5 cm
Htot
Inlet
Outlet
Filter
o'rings
L = 5 mm
d = 0.47 mm
d = 8 mm
B PBS filtration
Htot = 15 cm
Htot = 20 cm
Htot = 30 cm
Calibration
Domain I
Domain II
Flow rate [mL min-1]
Time [min]
Exp.
Fit
Total height, Htot [cm]
C Diluted blood filtration
R² = 0.90
D Whole blood filtration

**Figure S1. Schematic of the GµF set-up and gravity flow characterization.** (Related to Figure 1.) (A) The total column height ($H_{tot}$), including the sample height (H), the length of tubes 1 and 2 ($L_1$ and $L_2$) and the height of the filtration cartridge (h) determines the pressure, and consequently the flow rate. Insert shows close-up of the cartridge section, with the filter location and the tubes connections through Luer Lock junctions at the inlet and outlet. (B, C and D) Examples of GµF curves representing the flow rate evolution over time during filtration of (B) 10 mL of PBS, (C) 10 mL of diluted blood (1/6, v/v in PBS) and (D) 5-8 mL of whole blood through 8 µm diameter pore filters for $H_{tot}$ = 15, 20 and 30 cm, and the average flow rate in domain II as a function of the total height of the GµF set-up obtained experimentally (red dots) and calculated (black diamonds). Error bars correspond to the standard deviation of three replicated experiments.

**Table S1. Filter characteristics and geometry of the GµF set up.** (Related to Figures 1 and 2.) For each filter used, the pore diameter, the membrane thickness, the total number of pores (8 mm diameter filters) and the porosity are provided. The tube length ($L_1$) and the corresponding total height ($H_{tot}$) used in this work within the GµF set-up to achieve flow rates of 0.1 and 0.5 mL min$^{-1}$ are mentioned for all pore sizes.

| **Filter pore diameter (µm)** | **8** | **10** | **12** | **15** | **20** | **28** |
|---|---|---|---|---|---|---|
| | | | **Filter characteristics** | | | |
| **Thickness (µm)** | 20 | 20 | 20 | 40 | 40 | 40 |
| **Total number of pores per filter** | 80424 | 80424 | 80424 | 55850 | 31416 | 16029 |
| **porosity (%)** | 8 | 12.6 | 18 | 19.6 | 19.6 | 19.6 |
| | | | **Flow rate: 0.1 mL min$^{-1}$** | | | |
| **$L_1$ (cm)** | 12 | 11 | 7 | 6 | 6 | 6 |
| **$H_{tot}$ (cm)** | 22 | 21 | 17 | 16 | 16 | 16 |
| | | | **Flow rate 0.5 mL min$^{-1}$** | | | |
| **$L_1$ (cm)** | 66 | 64 | 56 | 47 | 47 | 47 |
| **$H_{tot}$ (cm)** | 76 | 74 | 66 | 57 | 57 | 57 |

*Note: For the CRCLM patients and some EOC patients a new generation of 8-µm-pore and 15-µm-pore filters with a porosity of 40% were used with 402,120 and 113,980 micropores, respectively.*

#### Flow rate evolution over time

Flow rate evolution during GµF was monitored for PBS, diluted blood (1:6 (v/v) in PBS) and whole blood through 8 µm diameter pore filters. For each fluid, flow rate was measured using three different tube lengths ($L_1$ = 5, 10 and 20 cm), corresponding to $H_{tot}$ = 15, 20 and 30 cm (Figure S1). Before filtration, the inlet tube was clamped at the top of Tube 1, below the top reservoir, and 10 mL samples were poured in the reservoir. Filtration started as the clamp was removed. During filtration, at known time intervals, sample droplets were collected from the cartridge outlet in 1.5 mL Eppendorf tubes, then weighed, and knowing the density of each fluid, the instantaneous flow rate for each time interval was then determined.

For all tested fluids, the flow rate evolution over time exhibited two domains, defined by the change in slop on the curve. For PBS (Figure S1B), flow rate quickly decreased during the first seconds to minutes (Domain I), then decreased according to the diminution of the fluid height until the end of filtration (Domain II). For diluted and whole blood (Figure S1C and D), initial flow rates also quickly decreased in domain I. Then, in domain II, flow rates remained constant, fluctuating around a single value for a few hours. As expected, average flow rates in domain II were lower for shorter initial fluid columns. For instance, for diluted blood, flow rates were 0.04, 0.09 and 0.15 mL $min^{-1}$, for $H_{tot}$ = 15, 20, and 30 cm, respectively.

The quick decrease in flow rate observed in domain I for each fluid was attributed to the fall of the sample on the filter once Tube 1 is unclamped. For 10 mL of diluted blood, the volume filtered in domain I corresponded to less than 10% of the total sample volume filtered (*i.e.* ~0.17 mL of whole blood before dilution).

For all fluids and column heights, average flow rates were determined from the pseudo-steady state in domain II over three replicated experiments. The 2-3 final data points, when flow rate dropped to zero at the very end of filtration, were not considered. For PBS, experimental flow rates in domain II were consistent with theoretical flow rates determined by estimating the flow resistance in the GµF set-up (Figure S1B and Table S2). For diluted and whole blood, correlation coefficients of 0.9 (Figure S1C and D) confirm the linear nature of the relationship between average flow rate in domain II and total column height ($H_{tot}$), thus allowing one to determine the tube length to use in order to obtain a specific flow rate.

#### Flow resistance

The contribution of each section of the GµF set-up to the flow resistance was estimated (Table S2) using Equation (1) for cylindrical sections (junctions and tubes) and Equation (2) for the filter with multiple parallel pores.

$$R_{tube} = \frac{8 \times \mu \times L_{tube}}{\pi \times r_{tube}^4} \qquad \text{Equation (1)}$$

$$R_{filter} = \frac{8 \times \mu \times t}{N_p \times \pi \times r_{pore}^4} \qquad \text{Equation (2)}$$

where μ is the coefficient of dynamic viscosity, $L_{tube}$, and $r_{tube}$ are the length and the internal radius of the considered cylindrical section, respectively, t, is the thickness of the porous membrane, and the $r_{pore}$ is the pore radius.

**Table S2**. **Flow resistance in the GμF set-up** and the relative contribution of each section to the total flow resistance for $L_l$ = 5, 10, 15 and 20 cm. (Related to Figure 1.)

| | **$L_l$ = 5 cm $H_{tot}$ = 15 cm** | | **$L_l$ = 10 cm $H_{tot}$ = 20 cm** | | **$L_l$ = 15 cm $H_{tot}$ = 25 cm** | | **$L_l$ = 20 cm $H_{tot}$ = 30 cm** | |
|---|---|---|---|---|---|---|---|---|
| | **R (Pa s m$^{-3}$)** | **R (%)** | **R (Pa s m$^{-3}$)** | **R (%)** | **R (Pa s m$^{-3}$)** | **R (%)** | **R (Pa s m$^{-3}$)** | **R (%)** |
| Luer lock | $3.47\ 10^{7}$ | **0.11** | $3.47\ 10^{7}$ | **0.08** | $3.47\ 10^{7}$ | **0.07** | $3.47\ 10^{7}$ | **0.06** |
| Tube 1 | $8.97\ 10^{9}$ | **27.1** | $1.79\ 10^{10}$ | **42.7** | $2.6910^{10}$ | **52.8** | $3.5910^{10}$ | **59.9** |
| Top junction | $5.82\ 10^{9}$ | **17.6** | $5.82\ 10^{9}$ | **13.8** | $5.82\ 10^{9}$ | **11.4** | $5.82\ 10^{9}$ | **9.7** |
| Filter | $3.45\ 10^{9}$ | **10.4** | $3.45\ 10^{9}$ | **8.2** | $3.45\ 10^{9}$ | **6.8** | $3.45\ 10^{9}$ | **5.8** |
| Bottom jct. | $5.82\ 10^{9}$ | **17.6** | $5.82\ 10^{9}$ | **13.9** | $5.82\ 10^{9}$ | **11.4** | $5.82\ 10^{9}$ | **9.7** |
| Tube 2 | $8.97\ 10^{9}$ | **27.1** | $8.97\ 10^{9}$ | **21.4** | $8.97\ 10^{9}$ | **17.6** | $8.97\ 10^{9}$ | **15.0** |
| Total Resistance | $3.31\ 10^{10}$ | **100** | $4.2010^{10}$ | **100** | $5.10\ 10^{10}$ | **100** | $6.00\ 10^{10}$ | **100** |

Considering the total resistance calculated for the GμF set-up with various tube lengths, theoretical flow rates were determined for PBS (Figure S1B). For $L_l$ =15 and 20 cm, experimental flow rates in domain II were in the same range as the calculated values. Flow rates of 1.9 *vs.* 2.2 ml min$^{-1}$ and 2.8 *vs.* 2.4 mL min$^{-1}$ were obtained for experiment *vs.* theory, for $L_l$ =15 and 20 cm, respectively.

However, for shorter tube lengths ($L_l$ = 5 cm), experimental flow rates deviate from theory and were found smaller than predicted. The contribution of tube 1 to the total fluid resistance ($R_{tube\ 1}$) was predominant for $L_l \geq 10$ cm (~60% of the total GμF resistance, $R_{tot}$, for $L_l$ = 20 cm) while the contribution of the cartridge section, including the filters and junctions ($R_{cartridge} = R_{top\ jct.} + R_{filter} + R_{bot.\ Jct.}$), represented less than 30% of the total flow resistance. $R_{tube\ 1}$ decreases with $L_l$, resulting in the increase of the contribution of $R_{cartridge}$ in $R_{tot}$. For instance, the contribution of $R_{cartridge}$ increased from ~25% with $L_l$ =

20 cm, to > 45%, with $L_1$ = 5 cm. Resistance calculations, based on the set-up geometry were most likely underestimated, in particular for the cartridge section that includes several junctions between the cartridge parts, the joints and the filter, therefore explaining the deviation of experimental data from theory for shorter tube lengths.

#### Filter porosity

For all filters used in this study, the average flow rate in Domain II was determined as described previously for $L_1$ = 5, 10 and 20 cm (Figure S2). In all cases, flow rate increased linearly with the total column height, allowing one to select the appropriate tube length to achieve a specific flow rate. As expected, filters with higher porosity yielded higher flow rate for similar column height. Interestingly, filters with 15, 20 and 28 µm pores, with thickness of 20, 40 and 40 µm respectively, but all with a 19.6% porosity, yielded similar average flow rates, suggesting that the flow resistance due to the filter is mostly driven by the open surface of the filter in this dimension range. For the two flow rates used in this study, the tube length ($L_1$) and the corresponding total height ($H_{tot}$) are provided in Table S1 for all filters.

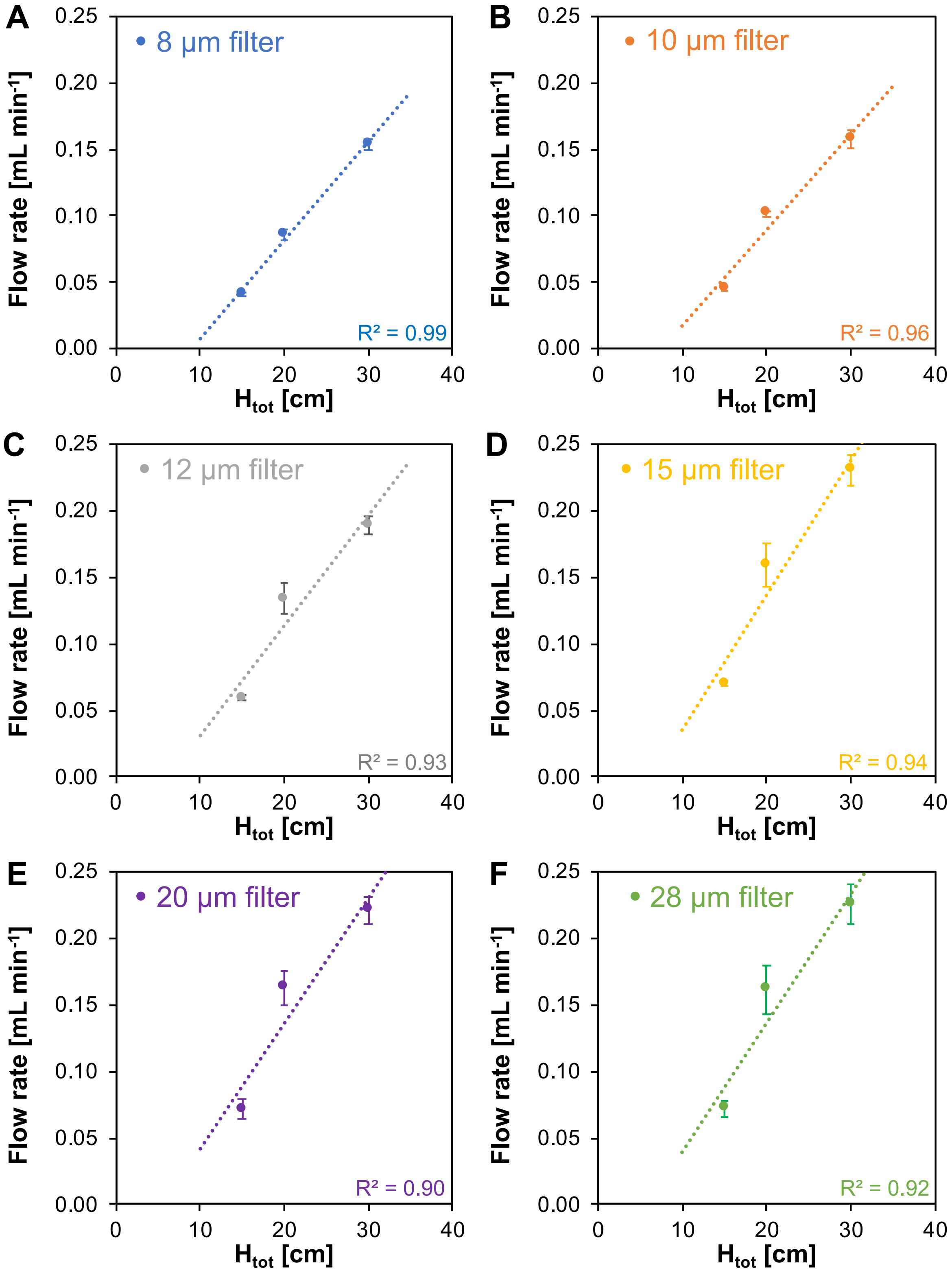

**Figure S2. GµF calibration for all pore sizes.** (Related to Figure 1.) Average flow rate in domain II for various column heights and for filters with pore diameter of (A) 8 µm, (B) 10 µm, (C) 12 µm, (D) 15 µm, (E) 20 µm, and (**F**) 28 µm. Error bars correspond to the standard deviation of three replicated experiments.

#### Transmembrane pressure and flow simulation

##### *Transmembrane pressure*

The microfilters used in this study consist of a porous membrane with multiple parallel cylindrical pores with diameter ranging from 8 to 28 µm. Porosity, thickness and total number of pores for each membrane filter are provided in Table S1. Considering a laminar flow through the multiple parallel openings, transmembrane pressure (△P) during filtration was determined using Equation (3), where Q is the flow rate, S is the membrane area, µ is the coefficient of dynamic viscosity, L is the membrane thickness, $N_p$ is the number of pores, and R is the pore radius (Table S3).

$$\Delta P = \frac{Q}{Np}\frac{128\,\mu\, l}{\pi D^4} \qquad \text{Equation (3)}$$

##### *Flow profile in pores*

Finite-element analysis comparing the flow profile during GµF and constant flow rate filtration (pump filtration) was performed using the COMSOL Multiphysics software. Flow velocity profiles were obtained by 3D simulations and flow speeds (ν) were determined using Equation (4), where R is the pore radius, µ is the coefficient of dynamic viscosity, and r is the distance from the center of the pore.

$$v = \frac{r^2 - R^2}{4\mu}\left(\frac{dp}{dx}\right) \qquad \text{Equation (4)}$$

Max velocity is when r = 0

$$v\ max = -\frac{R^2}{4\mu}\left(\frac{dp}{dx}\right)$$

$$V\ ave = \frac{1}{2}\ v\ max$$

$$V\ ave = -\frac{R^2}{8\mu}\left(\frac{dp}{dx}\right)$$

Maximum flow speeds ($\nu_{max}$) were determined for 8 µm and 15 µm filters for GµF and pump filtration. For GµF (constant pressure), ΔP was fixed at 5.2 Pa. Pump filtration was simulated with a constant flow rate of 0.1 mL $min^{-1}$. For both configuration, flow velocity profiles were simulated through a cell of 9 pores for 0% (9 open pores) and 22% clogging (same cell with 2/9 clogged pores) (Table S3 and Figure 2D). As expected, for GµF, clogging did not affect $\nu_{max}$, which remained ~830 µm $s^{-1}$, and for constant flow rate filtration, when clogging increased from 0% to 22%, a ~30% increase in $\nu_{max}$ was observed, yielding $\nu_{max}$ > 1mm $s^{-1}$. Average flow velocities ($\nu_{avg}$), determined using Equation (5) were 412 and 169 µm $s^{-1}$ for 8 µm and 15 µm filters, respectively.

$$\nu_{avg} = -\frac{R^2}{8 \times \mu} \times \frac{dP}{dx} \qquad \text{Equation (5)}$$

##### ***Shear Stress***

The frictional forces of the fluid acting on the cell surface during filtration is responsible for shear stress ($\tau$) that increases with fluid velocity and with viscosity. The maximum shear stress ($\tau_{max}$) in the pores was determined using Equation (7) for 8 µm and 15 µm filters (Table S3).

$$\frac{dv}{dr} = \frac{r}{2\mu}\left(\frac{dp}{dx}\right) \qquad \text{Equation (6)}$$

$$\tau = \mu\frac{dv}{dr} = \frac{r}{2}\left(\frac{dp}{dx}\right) \qquad \text{Equation (7)}$$

**Table S3. Flow characteristics in the GµF set-up.** (Related to Figure 2.) Transmembrane pressure (△P) calculated using Equation (3), for all filters and flow rates used in the study. Maximum flow speed ($\nu_{max}$) in each individual pore and of the maximum shear stress ($\tau_{max}$) along the walls of 8 µm and 15 µm filters used for scCTCs and cCTCs capture, respectively. 3D simulations were performed for constant pressure filtration (GµF) with △P = 5.75 Pa for 8 µm filter and 1.34 Pa for 15 µm filter, and for constant flow rate filtration (pump) at a flow rate of 0.1 mL $min^{-1}$.

| **Filter pore diameter (µm)** | **8** | | **10** | **12** | **15** | | **20** | **28** |
|---|---|---|---|---|---|---|---|---|
| | **Transmembrane pressure (Pa)** | | | | | | | |
| **Flow rate = 0.1 mL $min^{-1}$** | 5.75 | | 2.35 | 1.13 | 1.34 | | 0.75 | 0.38 |
| **Flow rate = 0.5 mL $min^{-1}$** | 28.73 | | 11.77 | 5.67 | 6.69 | | 3.77 | 1.92 |
| **Clogging %** | **0%** | **22%** | | | **0%** | **22%** | | |
| | **Max. speed in pores (µm $s^{-1}$)** | | | | | | | |
| **GµF** (△P = 5.75 Pa) | 826 | 826 | | | 338 | 338 | | |
| **Pump** (0.1 mL $min^{-1}$) | 826 | 1058 | | | 338 | 434 | | |
| | **Maximum shear stress (Pa)** | | | | | | | |
| **GµF** (△P = 5.75 Pa for 8 µm; 1.34 Pa for 15 µm) | 0.58 | 0.58 | | | 0.13 | 0.13 | | |
| **Pump** (0.1 mL $min^{-1}$) | 0.58 | 0.74 | | | 0.13 | 0.16 | | |

### B. Cluster area and cell diameter

#### Measure of cluster area

Using GµF, CTC capture relies on size, and due to the known heterogeneity in cell dimensions, clusters captured were characterized by measuring their surface area, therefore facilitating comparisons between experiments. The cluster area parameter was defined as the surface covered by a cluster on the filter after filtration. Cluster area was measured on bright field images using ImageJ software (Rasband, 2012), and an equivalent diameter ($Ø_{eq}$) was calculated considering clusters as perfect spheres (Figure S3A). $Ø_{eq}$ does not represent the cluster geometry and is used only for size range estimation and comparison.

#### Cell size in small clusters

Blood samples diluted 1:6 (v/v) in PBS and spiked with ~150-500 OV-90 clusters were filtered successively through all filters by order of decreasing pore size (28, 20, 15, 12, 10, then 8 µm diameter). Small clusters were captured on all filters and larger clusters (from 6-100+ cells) were mainly found on the first filters with the largest pores (28, 20, and 15 µm) (Figure 3C). For all filters, the diameter of each individual cell in small clusters (up to 5-6 cells) was measured (Figure S3B) and the diameter of the biggest cell was found to increase with the pore size, suggesting that the passage of small clusters is mostly limited by the largest cell size and deformability.

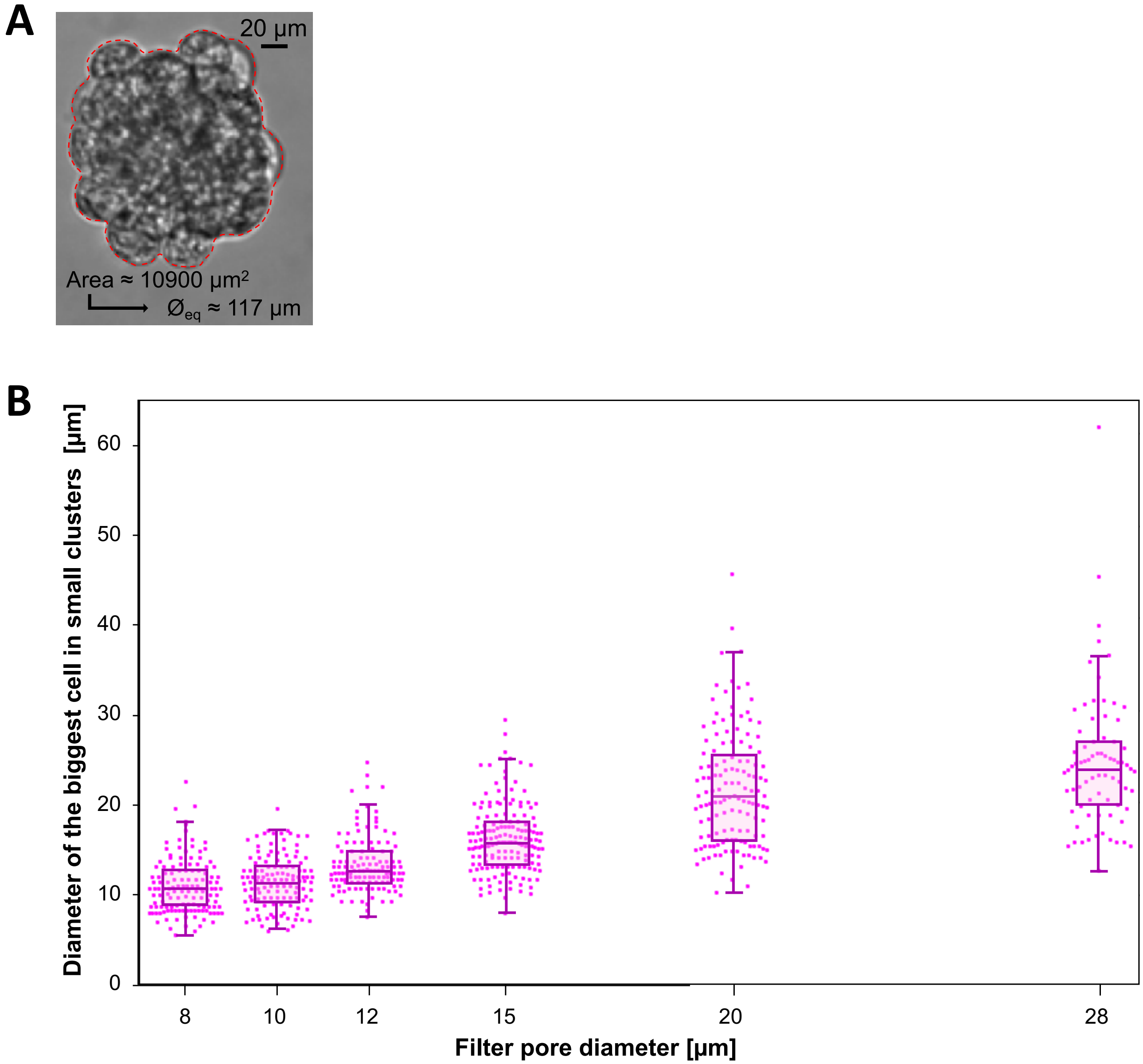

**Figure S3**. **Cluster area and cluster's cells diameter.** (Related to Figure 3.) (A) Example of the measure of the cluster area using ImageJ software on bright field images. Considering clusters as perfect spheres, an equivalent diameter ($Ø_{eq}$) is determined for comparison. (B) After successive filtrations through all filters by order of decreasing porosity (28, 20, 15, 12, 10, then 8 µm diameter), the diameter of the biggest cell in small clusters (< 6 cells) captured on each filter was measured (scatterplots), for three replicated experiments (n = 145, 300, and 550 clusters, respectively). The boxes contain values within the 25$^{th}$ and 75$^{th}$ percentiles, the whiskers correspond to the 91$^{st}$ and 9$^{th}$ percentiles, and the horizontal lines represent the medians. The biggest cell within small clusters limits their passage through pores.

### C. Growth of OV-90 and OVCAR-3 clusters captured from mouse blood

#### Migration assay

OV-90 and OVCAR-3-GFP clusters, captured from the blood of ovarian orthotopic mouse models, were released from the filters and maintained in culture as adherent cells. Once confluent, OV-90 and OVCAR-3 cells were harvested and migration assays were performed to characterize their growth in adherent layers (Figure S4A-D). About $5 \times 10^5$ OVCAR-3-GFP cells or OV-90 cells per milliliter were seeded in the two wells of a 2-well silicon insert (Ibidi, Germany) placed at the bottom of a Petri dish (well-well distance = 500 ± 50 µm). After overnight incubation, the silicon inserts were removed and the closure of the cell-free area (% closure) was monitored over time for both cell types. For each group of cells, the cell-free area was imaged and averaged over 10 images at each time point. For each condition, measurements done right after the removal of the silicon insert were used to set the reference for 0% closure. The increase of cell coverage over time (% closure) was thus determined by comparison with the reference.

Similar behaviors were observed for OV-90 and OVCAR-3-GFP, where cell coverage increased over time and reached a total closure (~100%) after ~6 days. However, OV-90 cells were observed to grow faster at first, with ~45% reduction of the cell-free area (closure ~35 to 80%) from day 2 to day 3, while with OVCAR-3-GFP, only ~25% reduction of the cell-free area (closure ~35 to 60%) was observed from day 2 to day 3.

#### Growth in suspension

OV-90 and OVCAR-3 clusters, isolated from the blood of ovarian orthotopic mouse model, were released from filters and seeded in wells of an ultra-low adhesion plate. Cell suspensions were imaged using bright field microscopy right after seeding, and over a few days of incubation (Figure S4E and S4F). Cluster area was measured using ImageJ software. Cluster growth in suspension was estimated by averaging the area of ~400-500 clusters, measured over a few days in three different wells. The evolution of the cluster size distribution over time is presented as box and whisker plots in figure S4G.

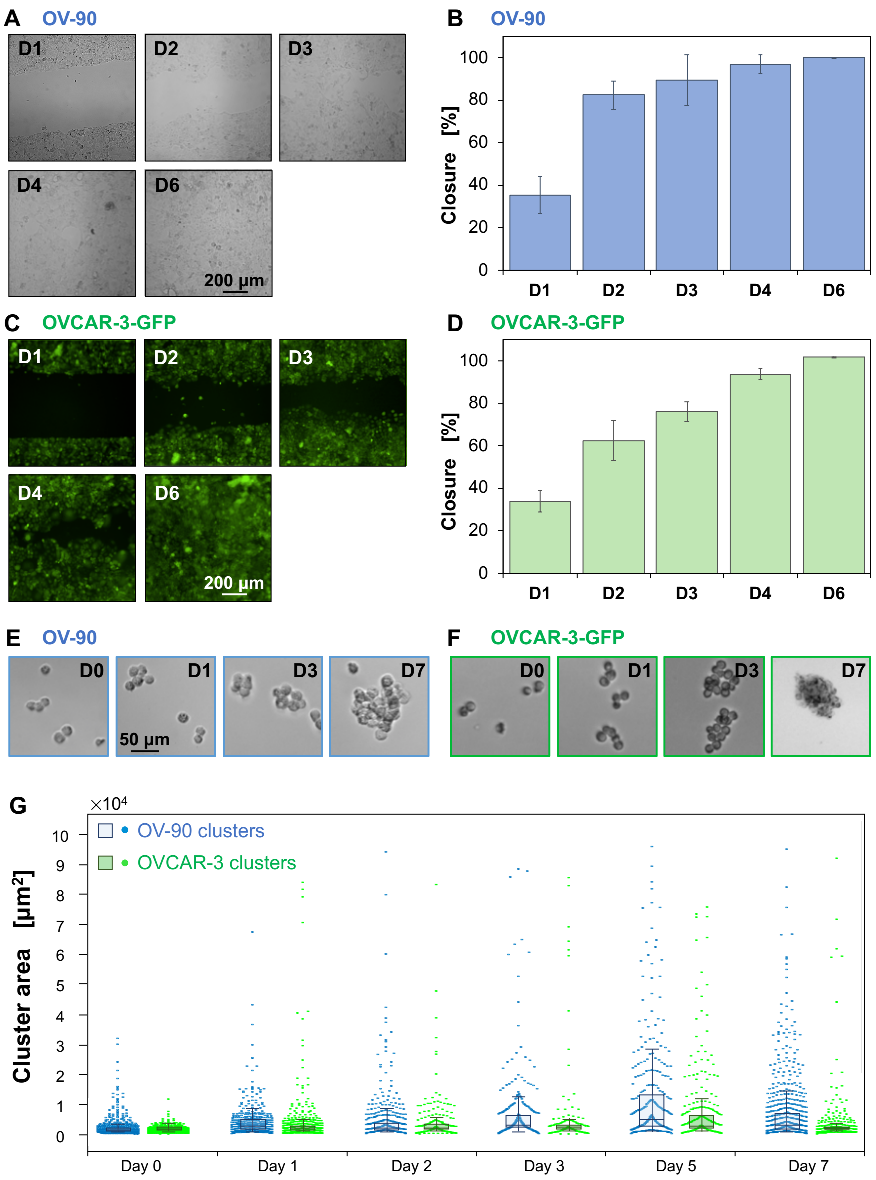

**Figure S4. Growth of OV-90 and OVCAR-3-GFP clusters previously isolated from mouse blood.** (Related to Figure 4.) (A-D) **Migration assays.** Representative images of (A) OV-90 (bright field) and (C) OVCAR-3-GFP (fluorescence) cluster growth overtime and closure of the cell-free area (% closure) for (B) OV-90 and (D) and OVCAR-3-GFP clusters respectively. (E-G) **OV-90 and OVCAR-3 cluster proliferation.** Bright field images of (E) OV-90 (top) and (F) OVCAR-3 (bottom) clusters from mouse blood, after capture and release in low-adhesion dishes. After 5-7 days, the morphology of OVCAR-3 vs. OV-90 clusters suggests that OVCAR-3 clusters are more sensitive to hypoxic conditions as cluster size increases (see main text). (G) OV-90 and OVCAR-3 cluster size distribution over time. The areas of 400-500 OV-90 (blue) and OVCAR-3 (green) clusters were measured over time after seeding in low-adhesion dishes, for three replicated experiments (scatterplots). The boxes contain values within the 25th and 75th percentiles, the whiskers correspond to the 91$^{st}$ and 9$^{th}$ percentiles, and the horizontal lines represent the medians.

### D. WBCs in cCTCs isolated from EOC patient samples

**Table S4. WBC count in cCTCs isolated from 10 EOC patient samples.**

| Patients | cCTCs | $WBC^+$ cCTCs | % $WBC^+$ cCTCs | Number of WBC in cCTC | Number of cells in $WBC^+$ cCTC |
|---|---|---|---|---|---|
| **OC1** | 186 | 0 | **0** | n/a | n/a |
| **OC2** | 13 | 0 | **0** | n/a | n/a |
| **OC3** | 9 | 1 | **11** | 2 | 4 |
| **OC4** | 4 | 2 | **50** | 1 | 3 |
| | | | | 1 | 3 |
| **OC5** | 20 | 5 | **25** | 1 | 3 |
| | | | | 1 | 3 |
| | | | | 1 | 3 |
| | | | | 1 | 3 |
| | | | | 2 | 4 |
| **OC11** | 178 | 0 | **0** | n/a | n/a |
| **OC12** | 98 | 9 | **9.2** | 1 | 3 |
| | | | | 1 | 3 |
| | | | | 1 | 3 |
| | | | | 1 | 3 |
| | | | | 1 | 3 |
| | | | | 1 | 5 |
| | | | | 3 | 5 |
| | | | | 1 | 3 |
| | | | | 1 | 3 |
| **OC13** | 31 | 1 | **3.2** | 1 | 5 |
| **OC14** | 11 | 0 | **0** | n/a | n/a |
| **OC15** | 1 | 0 | **0** | n/a | n/a |

### E. WBCs in cCTCs isolated from CRCLM patient samples

**Table S5. WBC count in cCTCs isolated from 13 CRCLM patient samples.**

| Patients | cCTCs | WBC+ cCTCs | % WBC+ cCTCs | Number of WBC in WBC+ cCTC | Number of cells in WBC+ cCTC |
|---|---|---|---|---|---|
| **CR1** | 6 | 0 | **0** | n/a | n/a |
| **CR2** | 4 | 0 | **0** | n/a | n/a |
| **CR3** | 27 | 10 | **37** | 1 | 2 |
| | | | | 3 | 3 |
| | | | | 2 | 4 |
| | | | | 2 | 3 |
| | | | | 3 | 7 |
| | | | | 1 | 2 |
| | | | | 2 | 4 |
| | | | | 2 | 4 |
| | | | | 1 | 2 |
| | | | | 2 | 5 |
| **CR4** | 37 | 8 | **22** | 3 | 7 |
| | | | | 1 | 2 |
| | | | | 1 | 3 |
| | | | | 1 | 5 |
| | | | | 1 | 4 |
| | | | | 1 | 2 |
| | | | | 1 | 4 |
| | | | | 1 | 2 |
| **CR5** | 9 | 2 | **22** | 2 | 8 |
| | | | | 1 | 2 |
| **CR6** | 22 | 0 | **0** | n/a | n/a |
| **CR7** | 5 | 0 | **0** | n/a | n/a |
| **CR8** | 4 | 0 | **0** | n/a | n/a |
| **CR9** | 10 | 4 | **40** | 1 | 3 |
| | | | | 1 | 3 |
| | | | | 1 | 2 |
| | | | | 4 | 5 |
| **CR10** | 16 | 0 | **0** | n/a | n/a |
| **CR11** | 77 | 27 | **35** | 2 | 3 |
| | | | | 1 | 3 |
| | | | | 2 | 4 |
| | | | | 2 | 5 |
| | | | | 1 | 3 |
| | | | | 1 | 3 |
| | | | | 4 | 6 |
| | | | | 2 | 3 |
| | | | | 2 | 4 |
| | | | | 2 | 3 |
| | | | | 3 | 5 |
| | | | | 1 | 3 |
| | | | | 2 | 4 |
| | | | | 1 | 2 |
| | | | | 2 | 4 |
| | | | | 2 | 3 |
| | | | | 3 | 4 |
| | | | | 1 | 3 |
| | | | | 2 | 3 |

| | | | | | |
|---|---|---|---|---|---|
| | | | | 1 | 2 |
| | | | | 1 | 2 |
| | | | | 1 | 3 |
| | | | | 1 | 2 |
| | | | | 2 | 3 |
| | | | | 1 | 2 |
| | | | | 1 | 2 |
| | | | | 2 | 8 |
| **CR12** | 74 | 9 | **12** | 1 | 3 |
| | | | | 1 | 3 |
| | | | | 1 | 4 |
| | | | | 1 | 3 |
| | | | | 1 | 2 |
| | | | | 2 | 6 |
| | | | | 1 | 2 |
| | | | | 1 | 2 |
| | | | | 1 | 2 |
| **CR13** | 45 | 15 | **33** | 1 | 3 |
| | | | | 1 | 2 |
| | | | | 1 | 5 |
| | | | | 2 | 5 |
| | | | | 1 | 4 |
| | | | | 2 | 3 |
| | | | | 4 | 10 |
| | | | | 3 | 5 |
| | | | | 3 | 4 |
| | | | | 3 | 5 |
| | | | | 1 | 2 |
| | | | | 1 | 3 |
| | | | | 3 | 6 |
| | | | | 1 | 5 |
| | | | | 2 | 4 |

### References

Rasband, W. S. (2012). ImageJ: image processing and analysis in Java. ASCL*, 1***,** 06013.
Ostadfar, A. in Biofluid Mechanics (ed Ali Ostadfar) 1-60 (Academic Press, 2016)
